# A comparison of dopaminergic and cholinergic populations reveals unique contributions of VTA dopamine neurons to short-term memory

**DOI:** 10.1101/2020.07.26.221713

**Authors:** Jung Yoon Choi, Heejae Jang, Sharon Ornelas, Weston Fleming, Daniel Fürth, Jennifer Au, Akhil Bandi, Ilana B. Witten

## Abstract

We systematically compared the contribution of two dopaminergic and two cholinergic ascending populations to a spatial short-term memory task in rats. In ventral tegmental area dopamine (VTA-DA) and nucleus basalis cholinergic (NB-ChAT) populations, trial-by-trial fluctuations in activity during the delay period related to performance with an inverted-U, despite the fact that both populations had low activity during that time. Transient manipulations revealed that only VTA-DA neurons, and not the other three populations we examined, contributed causally and selectively to short-term memory. This contribution was most significant during the delay period, when both increases or decreases in VTA-DA activity impaired short-term memory. Our results reveal a surprising dissociation between when VTA-DA neurons are most active and when they have the biggest causal contribution to short-term memory, while also providing new types of support for classic ideas about an inverted-U relationship between neuromodulation and cognition.

## Introduction

Short-term memory (Baddeley, 1986; Baddeley and Hitch, 1974; Erlich et al., 2011; Funahashi et al., 1993; Fuster and Alexander, 1971; Inagaki et al., 2019; Kamigaki and Dan, 2017; Kopec et al., 2015; Kubota and Niki, 1971; Liu et al., 2014; Miller et al., 2018; Romo et al., 1999) is a fundamental cognitive process with distinct temporal components: a “sample period” in which new information is updated into short-term memory, a “delay period” in which the memory is maintained, and ultimately a behavioral readout based on the memory (“choice period”). Although neuromodulators have been implicated in short-term memory (Brozoski et al., 1979; Clark and Noudoost, 2014; Croxson et al., 2011; Everitt and Robbins, 1997; Hasselmo and Stern, 2006; Ott and Nieder, 2019; Sun et al., 2017), it remains unclear which neuromodulators are most relevant, and which temporal component of short-term memory they support.

For example, DA has been implicated in short-term memory through pioneering experiments that pharmacologically manipulated DA receptors in PFC in monkeys performing short-term memory tasks (Arnsten et al., 1994; Cai and Arnsten, 1997; Floresco and Phillips, 2001; Murphy et al., 1996; Sawaguchi and Goldman-Rakic, 1991; Vijayraghavan et al., 2007; Williams and Goldman-Rakic, 1995; Zahrt et al., 1997). This work suggested that DA has an “inverted-U” influence on short-term memory and on memory-related activity during the delay period. In other words, too much or too little DA is detrimental to short-term memory, while intermediate levels enhance short-term memory. From these experiments, the idea arose that optimal levels of DA in prefrontal cortex (PFC) during the delay period serves to stabilize memory-related activity (Figure 1a; Arnsten, 1997; Arnsten et al., 2012; Cools and D’Esposito, 2011; Gibbs and D’Esposito, 2005).

**Figure 1.**
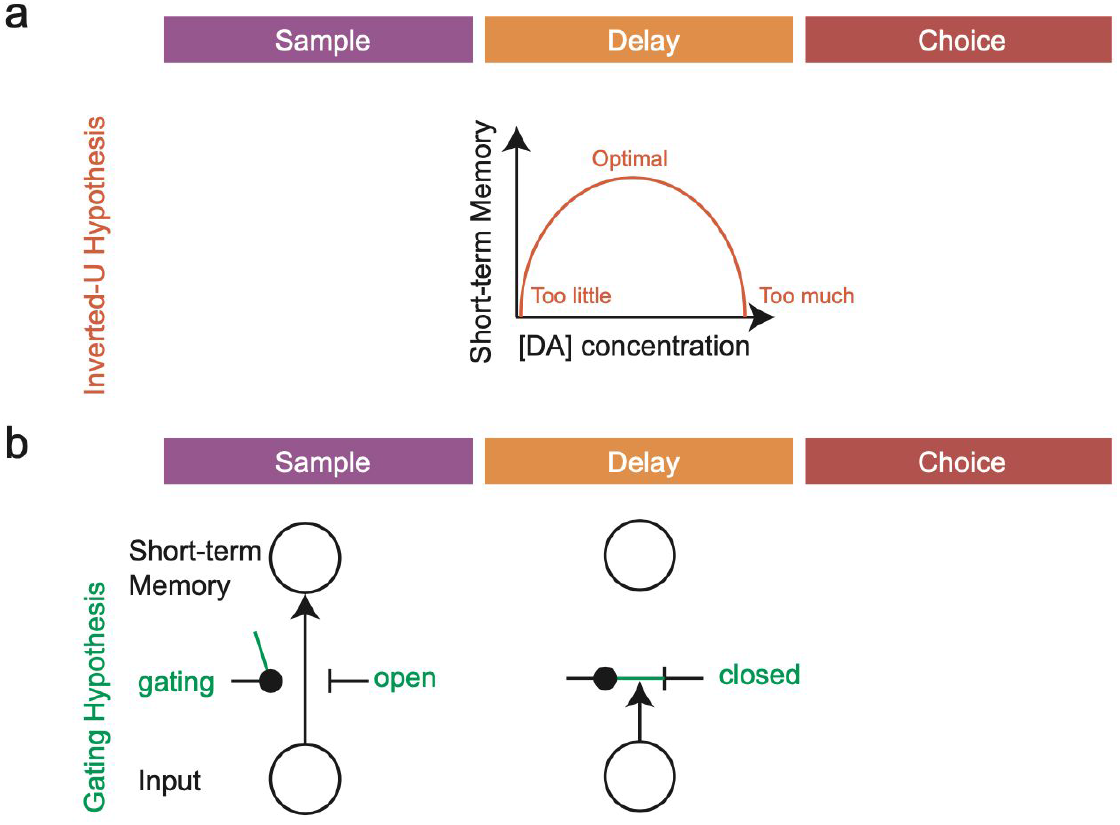
Gating and inverted-U hypotheses emphasize the contribution of DA to short-term memory in the sample and delay period, respectively. (a) Schematic of inverted-U hypothesis adapted from(Cools and Robbins, 2004). In this framework, DA contributes to maintaining a memory item during the delay period. (b) Schematic of gating hypothesis, adapted from(Hazy et al., 2007). In this framework, DA contributes to updating of new information during the sample period.

However, integrating these findings with the understanding that has emerged based on direct recordings of activity in DA neurons has presented a challenge to this idea. DA neurons with cell bodies in the ventral tegmental area (VTA) and substantia nigra (SNc) send projections to the striatum, PFC, and other forebrain regions. These neurons, which are thought to provide the major source of DA to their forebrain targets, are known to respond transiently to unexpected rewards and reward-predicting cues (Bayer and Glimcher, 2005; Cohen et al., 2012; Ellwood et al., 2017; Ljungberg et al., 1991; Parker et al., 2016; Roesch et al., 2007; Schultz, 1986, 1998; Schultz et al., 1993). This signal has been interpreted as a reward prediction error, which is thought to support reinforcement learning (Chang et al., 2016; Parker et al., 2016; Steinberg et al., 2013). On the other hand, dopamine neurons are not known to be active during the delay period of tasks with short-term memory components, when rewards and reward-predicting cues are absent (Cohen et al., 2012; Ljungberg et al., 1991; Matsumoto and Takada, 2013).

Thus, the “gating” theory of short-term memory has been proposed to integrate the role of DA in encoding a reward prediction error signal, with the idea that it regulates short-term memory (Figure 1b; Braver and Cohen, 1999, 2000; O’Reilly and Frank, 2006). In this model, phasic bursts of DA at the times of reward-predicting events serve to open the “gate” of short-term memory, and update relevant items into short-term memory. Low levels of DA during the delay period allow the gate to remain closed and prevent distractors from overwriting the memory item.

In particular, the gating theory suggests that phasic DA at the time of *updating* is critical to short-term memory, while the classic ideas based on pharmacology suggest that tonic levels of DA during the *delay* period are more important (Figure 1). In order to directly test these two ideas, we must understand *when* DA contributes to short-term memory – does DA affect the updating of short-term memory with new information during the sample period, or is it more important during the delay period?

Addressing this question requires knowing which DA subpopulations are relevant to short-term memory, so that we can selectively record from and manipulate the relevant neurons with appropriate temporal resolution. The two major ascending sources of DA to the forebrain arise from the VTA and SNc. Although VTA-DA sends stronger projections to medial prefrontal cortex than SNc (Beckstead et al., 1979; Lindvall et al., 1978), there is evidence that PFC receives input from both subpopulations (Williams and Goldman-Rakic, 1998). In addition, previous papers have suggested a role for SNc (or dorsal striatum DA, the major target of SNc) in short-term memory (e.g. Bellissimo et al., 2004; Landau et al., 2009; Matsumoto and Takada, 2013).

In addition to determining which DA neurons are relevant to short-term memory, we also wanted to know if the contribution of DA to short-term memory is unique relative to other neuromodulators. We chose to focus on ascending cholinergic (ChAT) neurons arising from the basal forebrain regions –nucleus basalis (NB) or medial septum (MS)– given previous work implicating these populations in short-term memory and other cognitive processes (Croxson et al., 2011; Hasselmo, 2006; Hasselmo and Sarter, 2011; Hasselmo and Stern, 2006; Sun et al., 2017).

Thus, we employed fiber photometry and optogenetics to monitor and manipulate DA and ChAT neurons with sub-second resolution in rats performing a spatial short-term memory task in an operant chamber. We found that DA neurons in the VTA and SNc, as well as ChAT neurons in the NB, encoded task events more than the animal’s movement in the chamber. These task-encoding populations had elevated activity during the sample, choice and reward periods. Instead, during the delay period, VTA-DA had low activity, consistent with the gating theory. Interestingly, during the delay period, the natural pattern of activity in this population had an inverted-U relationship with performance. Only DA neurons in the VTA, and not the other populations, causally and specifically contributed to task performance. In particular, VTA-DA inhibition during the sample period led to impairments in short-term memory, providing some causal support for the gating theory. In addition, despite the low activity during the delay period, both optogenetic increases and decreases of VTA-DA activity during that period lead to impairment in performance, providing the first causal support (to our knowledge) at the level of VTA-DA neuron firing for the inverted-U hypothesis. Together, this work identifies a unique role for VTA-DA neurons in short-term memory, and provides new correlational and causal support for both the gating theory and the inverted-U hypothesis, implying that VTA-DA contributes to both the updating and the maintenance of short-term memory.

## Results

### Rats performed a delayed nonmatch to position task (DNMTP) task during optical recording of DA and ChAT neurons

Rats were trained on a rodent spatial short-term memory task known as delayed non-match to position (Figure 2a; DNMTP; Akhlaghpour et al., 2016; Dunnett et al., 1988). In the DNMTP task, rats are presented with a sample lever in one of two possible locations on the front wall of the chamber (“sample presentation”). Upon pressing the lever (“sample press”), the lever retracts and the nosepoke on the back wall of the chamber is illuminated. The rat then initiates the delay period by entering the nose poke (“delay start”). After a delay of either 1, 5, or 10 s, when the rat re-enters the nose poke, both levers are presented on the front wall (“choice presentation”). To obtain a water reward, the rat must press the lever that does not match the initial sample lever (“choice press”). Trained rats performed well above chance and displayed a delay-dependent decline in performance (Figure 2b; one-way ANOVA, accuracy explained by delay; p < 0.001 for delay; n = 34 rats).

**Figure 2.**
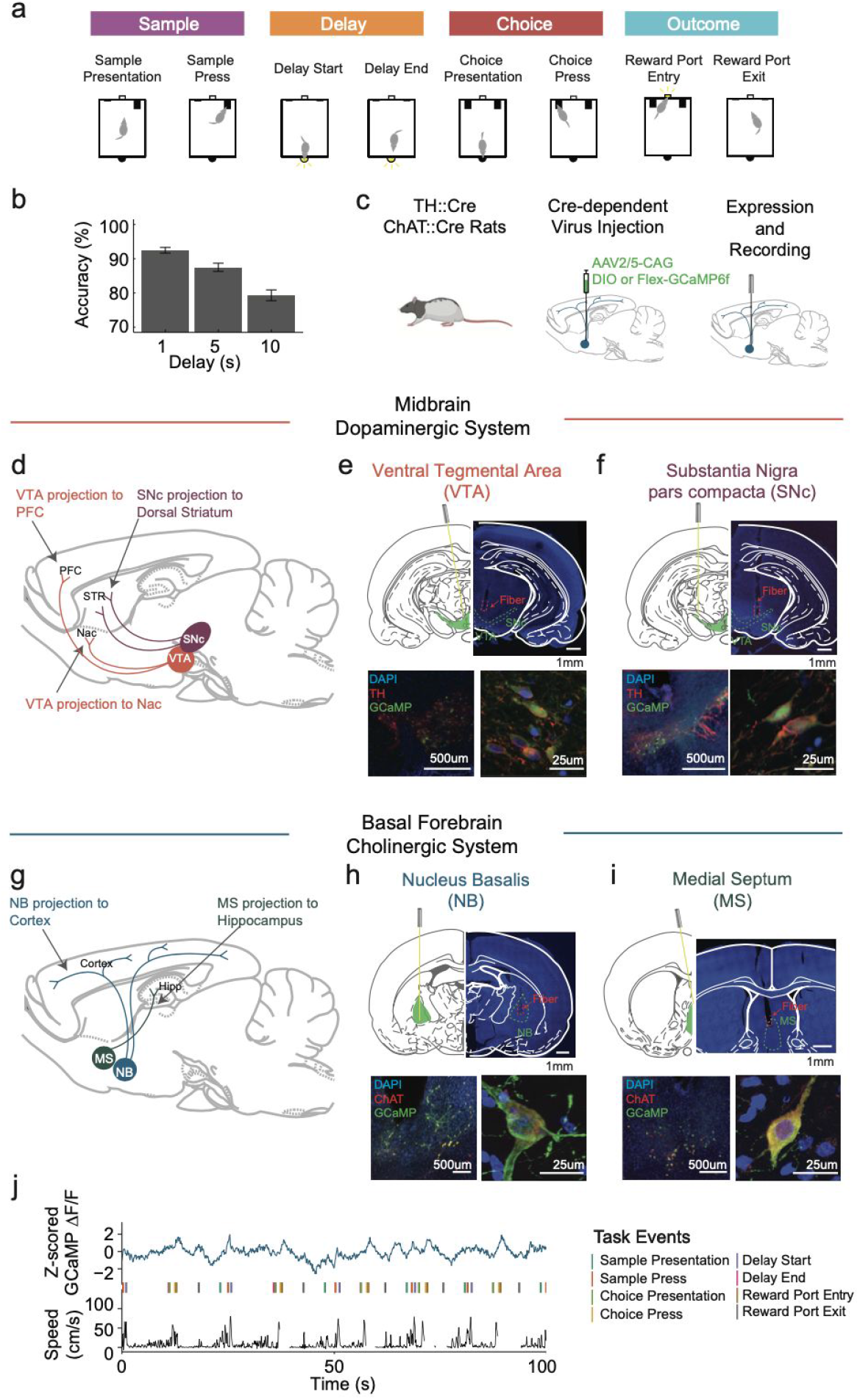
Fiber photometry recordings of VTA-DA, SNc-DA, NB-ChAT and MS-ChAT neurons in rats performing a spatial short-term memory task. (a) Schematic of the delayed non-match to position task (DNMTP). The sample period starts with the sample lever presentation on either the left or right side of the chamber (“Sample Presentation”). Pressing the sample lever (“Sample Press”) triggers the nosepoke on the back of the chamber to be illuminated. The delay period initiates when the rat makes a nosepoke, which turns off the nosepoke light (“Delay Start”). After the delay period (1, 5, or 10s), the nosepoke is again illuminated, signalling that the delay period is over (“Delay End”). Upon making another nosepoke, choice levers are presented (“Choice Presentation”). The rat needed to press the “non-match” lever (“Choice Press”) to be rewarded with a drop of water during the outcome period. Colored bars delineate the duration of the sample, delay, choice, and outcome periods relative to the task events. Note that choice period denotes the choice readout period, as opposed to when the choice is necessarily being made. (b) Behavioral performance of trained rats during fiber photometry recordings (performance during recordings from n=34 rats; bar and errorbar represent mean + sem across rats). Rats performed well above chance (50%), and showed delay-dependent impairment in accuracy (one-way ANOVA, accuracy explained by delay; p<0.001 for delay; n=34 rats). (c) Cell type-specific expression of GCaMP6f was obtained using TH::Cre or ChAT::Cre rats and Cre-dependent GCaMP virus (AAV2/5-CAG-DIO or FLEX-GCaMP6f) (d) Schematic of midbrain DA system. Two nuclei of interest are DA neurons the Ventral Tegmental Area (VTA-DA) and Substantia Nigra pars compacta (SNc-DA). (e-f) GCaMP6f (green) is specifically expressed in DA neurons (red) in the VTA (e) and SNc (f). (g) Schematic of basal forebrain ChAT system. Two nuclei of interest are ChAT neurons in the Nucleus Basalis (NB-ChAT) and Medial Septum (MS-ChAT). (h-i) GCaMP6f (green) is specifically expressed in ChAT neurons (red) in the NB (h) and MS (i). (j) Example recording trace showing simultaneous acquisition of time-varying GCaMP6f fluorescence from NB-ChAT neurons (top), timestamps of each task event (middle), and the rat’s speed in the chamber (bottom).

After training, rats were injected with a Cre-dependent AAV2/5 GCaMP6f virus in the VTA or the SNc in the case of TH::Cre rats (Figure 2d-f) or in the NB or MS in the case of ChAT::Cre rats (Figure 2g-i), and implanted with an optical fiber at the same location for fiber photometry recordings (Figure 2c; Supplementary Figure 1). We recorded time-varying GCaMP fluorescence during the task, along with the animal’s head position in the chamber and the timestamps for each task event (Figure 2j).

### VTA-DA, SNc-DA, and NB-ChAT populations (but not MS-ChAT) primarily encode task events rather than the rats’ speed

Before examining in detail the neural correlates of behavioral events, we determined if the animals’ movement in the chamber provided a better explanation of neural activity than the events themselves. This is a possible confound in interpreting neural correlates of events, given that in an operant task, an animal’s movement may correlate with the timing of task events, and therefore apparent neural correlates of task events may be better explained as neural correlates of movement (eg. speed may be lower during the delay or reward period, or higher before the delay period, when the animal traverses the chamber).

Thus, we compared the predictive power of linear encoding models (Engelhard et al., 2019; Lovett-Barron et al., 2019; Musall et al., 2019), in which the GCaMP signal was predicted based on different sets of predictors: either only speed (“speed model”), only task events (“event model”), or a full model based on both task events and speed (“event and speed model”, model schematic in Figure 3a, see Methods for details on encoding models). This revealed that the time-varying GCaMP signal in VTA-DA, SNc-DA, and NB-ChAT was better explained by the task events than speed, whereas speed explained GCaMP in MS-ChAT as well as task events (Figure 3b; one-way ANOVA, R^2^ over 3-fold cross validation for different encoding models: p<0.001 for VTA-DA, p<0.001 for SNc-DA, p<0.001 for NB-ChAT, p<0.001 for MS-ChAT; post-hoc pairwise t-test for difference between speed model and task events model, with bonferroni correction for multiple comparisons: p<0.001 for VTA-DA, p<0.001 for SNc-DA, p <0.001 for NB=ChAT, p=0.86 for MS-ChAT; n=10 recording sites for VTA-DA, n=13 for SNc-DA, n=18 NB-ChAT, n=7 for MS-ChAT; Supplementary Figure 2 for the speed encoding of MS-ChAT; Supplementary Figure 3 for the visualization of task event kernels learned from the full model).

**Figure 3.**
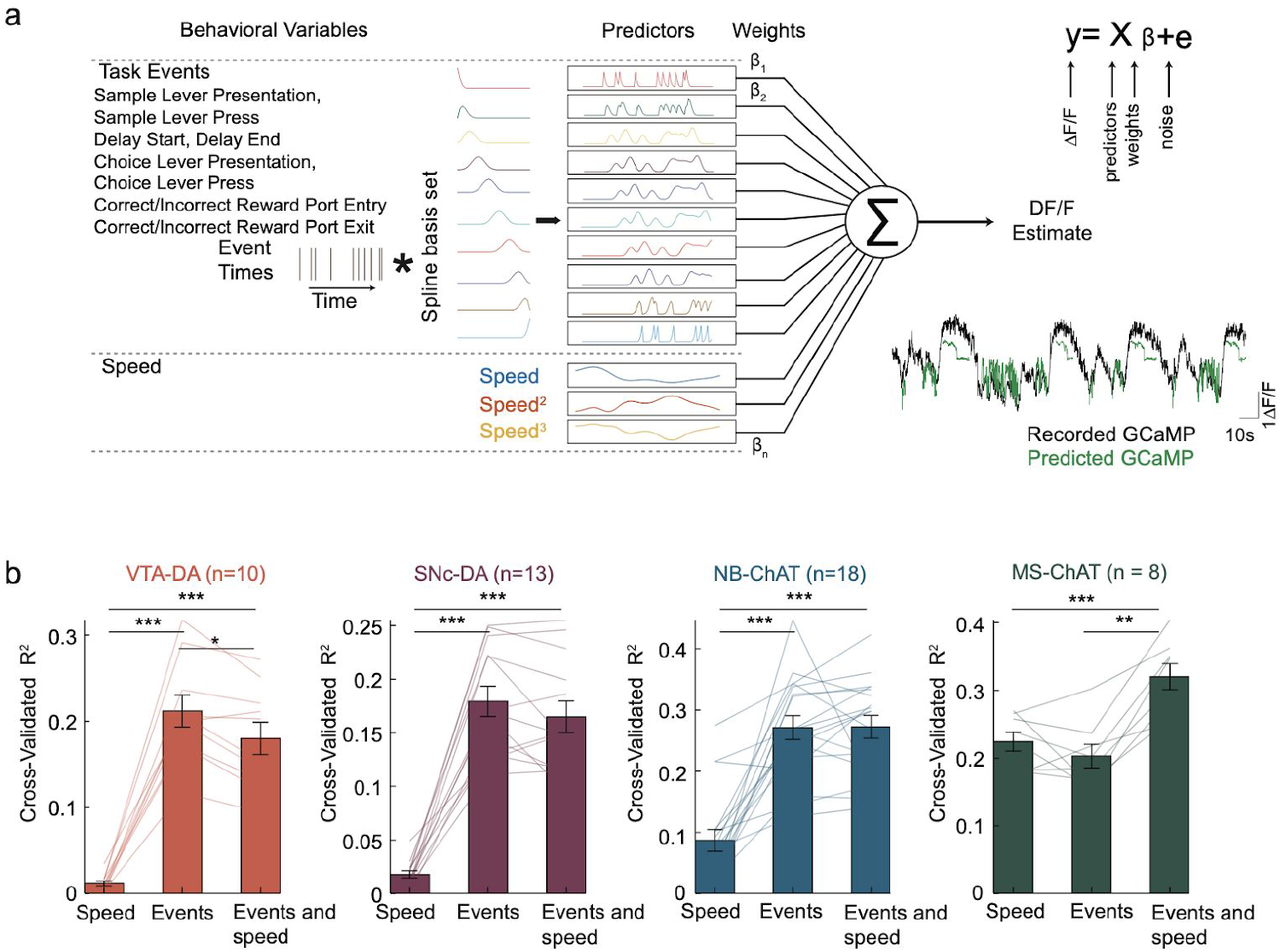
VTA-DA, SNc-DA, and NB-ChAT, but not MS-ChAT, better encode task events than speed. (a) Schematic of the full encoding model, in which GCaMP at each timepoint was predicted based on both task events and the rat’s speed. For each task event, a set of 10 predictors was created by convolving that task event’s time stamps with a spline basis set, in order to allow temporally delayed versions of each event to predict GCaMP fluorescence. Speed predictors include first, second, and third-degree polynomials of the animal’s speed at each point in time. (b) Three encoding models (x-axis) were generated and compared on held-out data: 1) a model with only speed predictors, 2) a model with only task event predictors, 3) the full model with both task event and speed predictors. In the VTA-DA, SNc-DA, and NB-ChAT populations, the task events model performed better than the speed model, while in the MS-ChAT population, the speed model and the task events model were comparable (one-way ANOVA, R2 explained by each encoding model: p<0.001 for VTA-DA, p<0.001 for SNc-DA, p<0.001 for NB-ChAT, p<0.001 for MS-ChAT; post-hoc pairwise t-test comparing difference between speed model and task events model, with bonferroni correction for multiple comparisons: p<0.001 for VTA-DA, p<0.001 for SNc-DA, p <0.001 for NB-ChAT, p=0.86 for MS-ChAT; n=10 recording sites for VTA-DA, n=13 for SNc-DA, n=18 for NB-ChAT, or n=8 for MS-ChAT). Bars and error bars indicate mean + sem across recording sites. Each dot represents a recording site. R2 for each recording site was obtained by averaging over 3-fold cross-validations.

### VTA-DA, SNc-DA and NB-ChAT neurons have elevated activity during the sample, choice, and outcome periods, but not the delay period

Given that task events were good predictors of the variance of GCaMP fluorescence in VTA-DA, SNc-DA, and NB-ChAT neurons, we further examined how neural activity correlated with each event in those populations by time-locking the GCaMP signal to each event (Figure 4a-i; Supplementary Figure 4). We observed some commonalities in the activity profiles across these task-encoding populations. For example, transient elevation of GCaMP fluorescence in relation to task events was evident across the sample, choice and reward period in all three populations (Figure 4j, 4l, 4n; one-way ANOVA, average GCaMP explained by sample, delay, choice or outcome epoch; p<0.001 for VTA-DA, p<0.003 for SNc-DA, p=0.002 for NB-ChAT; n=10 recording sites for VTA-DA, n=13 for SNc-DA, n=18 for NB-ChAT).

**Figure 4.**
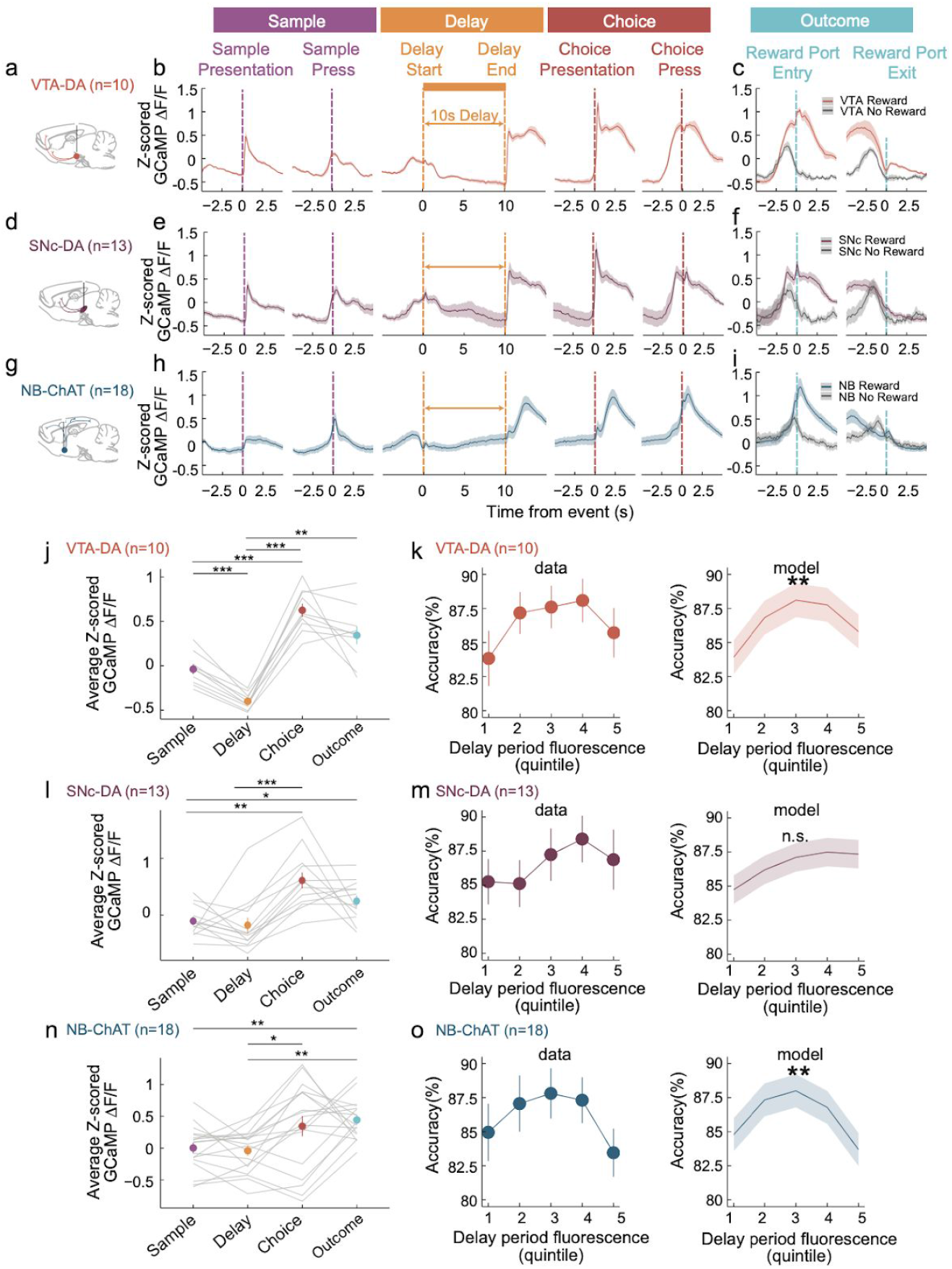
During the delay period, GCaMP fluorescence in VTA-DA and NB-ChAT relates to performance with an inverted-U, despite relatively low fluorescence at that time. (a) Schematic of fiber photometry recordings from VTA-DA. (b) Z-scored GCaMP fluorescence from VTA-DA recordings, time-locked to each task event during the sample, delay and choice periods (mean+sem across recordings, n=10 recording sites). Data from all 10s delay trials. (c) Z-scored GCaMP fluorescence from VTA-DA recordings during the outcome period, separated by rewarded and unrewarded trials (mean+sem across recording sites, n=10 recording sites). Data from all 10s delay trials. (d-f) Same as (a-c) but for SNc-DA recordings (n=13 recording sites). (g-i) Same as (a-c) but for NB-ChAT recordings (n=18 recording sites). (j) Average Z-scored GCaMP fluorescence during the sample, delay, choice, and outcome periods from VTA-DA recordings (10s delay trials). Each dot and errorbar represents mean +sem across recordings (n=10 recording sites). Average GCaMP activity was significantly lower in the delay period compared to the sample, choice, or outcome periods. (post-hoc pairwise t-test with bonferroni correction for multiple comparisons; p<0.001 for the difference between delay and sample, p<0.001 for the difference between delay and choice, p=0.001 for the difference between delay and outcome; n=10 VTA-DA recording sites). (k) Left: Accuracy relative to delay period fluorescence in VTA-DA. To relate delay period fluorescence to accuracy, we ranked all trials according to their average delay period fluorescence for each recording site (n=10 recording sites) and delay period duration (n=3 delay durations). For trials in each quintile for each delay period duration, we plotted the average accuracy versus the fluorescence quintile, averaging across delay period duration and then recording sites (mean+sem across recording sites, n=10 recording sites). Note that calculating GCaMP fluorescence quintiles separately for each delay accounted for delay-dependent differences in fluorescence and allowed visualization of the delay-independent relationship between fluorescence and accuracy. Right: Accuracy relative to delay period fluorescence predicted from the model fit to the data on the left (mixed-effect linear regression, accuracy predicted based on first and second degree polynomial of delay period fluorescence quintile, delay, and random effect of individual recording site; p=0.005 for the second degree polynomial; mean+sem across recording sites, n=10 recording sites). See Supplementary Figure 10g for additional statistical analyses of the inverted-U relationship. (l) same as (j) but for SNc-DA recordings (n=13 recording sites). Average GCaMP fluorescence was significantly lower in the delay period compared to the choice period, but not different from the sample nor outcome periods (post-hoc pairwise t-test with bonferroni correction for multiple comparisons; p=1.00 for the difference between delay and sample, p<0.001 for the difference between delay and choice, p=0.28 for the difference between delay and outcome; n=13 SNc-DA recording sites). (m) same as (k) but for SNc-DA recording sites (mixed-effect linear regression, accuracy predicted based on first and second degree polynomial of delay period fluorescence quintile, delay, and random effect of individual recording site; p=0.45 for the second degree polynomial, n=13 recording sites). (n) same as (j) but from NB-ChAT recordings (n=18 recording sites). Average GCaMP activity was significantly lower in the delay period compared to the choice and outcome periods, but not different from the sample periods (post-hoc pairwise t-test with bonferroni correction for multiple comparisons; p=1.00 for the difference between delay and sample, p=0.03 for the difference between delay and choice, p=0.002 for the difference between delay and outcome; n=18 NB-ChAT recording sites). (o) same as (k) but in NB-ChAT recording sites (mixed-effect linear regression, accuracy predicted based on first and second degree polynomial of delay period fluorescence quintile, delay duration, and random effect of individual recording site; p = 0.008 for the second degree polynomial, n=18 recording sites)

We did not observe elevation of GCaMP fluorescence during the delay period in any population. In VTA-DA, fluorescence during the delay period was significantly lower than during the sample or choice periods (Figure 4j; post-hoc pairwise t-test with bonferroni correction for multiple comparisons; p<0.001 for the difference between delay and sample, p<0.001 for the difference between delay and choice, n=10 VTA-DA recording sites). In SNc-DA and NB-ChAT recordings, the delay period fluorescence was not significantly different from the sample period, but significantly lower than the choice period (Figure 4l, 4n; post-hoc pairwise t-test with bonferroni correction for multiple comparisons; p=1.00 for SNc-DA, p=1.00 for NB-ChAT for the difference between delay and sample; p<0.001 for SNc-DA, p=0.03 for NB-ChAT for the difference between delay and choice; n= 13 SNc-DA, 18 NB-ChAT recording sites).

Finally, in both VTA-DA and SNc-DA populations, activity was higher during the choice period relative to the sample (Figure 4j, 4l; post-hoc pairwise t-test with bonferroni correction for multiple comparisons; p<0.001 for VTA-DA, p=0.002 for SNc-DA for the difference between sample and choice; n=10 VTA-DA, 8 SNc-DA recording sites). The higher activity during the choice period can be interpreted as modulation by a temporally discounted reward expectation function (Fiorillo et al., 2008; Kobayashi and Schultz, 2008; Mazur, 1987; Richards et al., 1997; Roesch et al., 2007; Samuelson, 1937; Starkweather et al., 2017), and therefore consistent with reward prediction error. This is because the sample and the choice periods involve the same stimulus and action, but the choice period was more proximal to the reward than the sample period. Relatedly, choice period activity was negatively correlated with delay duration in VTA-DA and SNc-DA populations, consistent with reward prediction error in that shorter delays reflect an earlier-than-expected outcome (Supplementary Figure 5).

In comparison, the NB-ChAT population choice period activity was not significantly higher relative to the sample (Figure 4n; post-hoc pairwise t-test with bonferroni correction for multiple comparisons; p=0.09 for the difference between sample and choice; n=10 NB-ChAT recording sites) and uncorrelated with delay duration (Supplementary Figure 5). Another distinction between NB-ChAT and the DA populations was that NB-ChAT preferentially responded to lever press action whereas VTA-DA preferentially responded to lever presentation cue (while SNc-DA had mixed selectivity for cue and action; Supplementary Figure 6).

As a control, we recorded from animals in which GFP and not GCaMP was expressed. In that case, we did not observe a similar pattern of modulation of fluorescence relative to task events (Supplementary Figure 7).

Additionally, we examined the spatial distribution of response selectivity within each region based on a careful anatomical reconstruction of recording fiber placement. Pairwise correlations between recording sites revealed that GCaMP responses were highly homogeneous in VTA-DA and MS-ChAT populations, and heterogeneous in SNc-DA and NB-ChAT populations (Supplementary Figure 8). Interestingly, NB-ChAT responses were spatially organized along the medio-lateral axis (Supplementary Figure 8). Furthermore, the medial and lateral subregions of NB-ChAT received topographic input from the medial and lateral subregions of the striatum, respectively (Supplementary Figure 9).

In summary, neural correlates in VTA-DA neurons during this task were consistent with the gating theory (Figure 1b), in that there was elevated activity during the sample period, and suppressed activity during the delay period. Activity was further elevated during the choice period, which can be considered as consistent with reward prediction error, and therefore the gating theory.

### During the delay period, VTA-DA and NB-ChAT activity relates to performance with an inverted-U relationship

Although delay period activity was relatively low in VTA-DA and NB-ChAT, we found an interesting relationship between activity during the delay period and task accuracy in both populations. Specifically, the average task accuracy as a function of delay period fluorescence followed an inverted-U relationship (Figure 4k,m,o). Statistically, this was confirmed with a mixed-effect linear regression in which accuracy was predicted based on the first and second degree polynomial of delay period fluorescence quintile (as well as delay period duration and a random effect of individual recording site). For both VTA-DA and NB-ChAT, but not SNc-DA, the second degree polynomial of delay period fluorescence was statistically significant, indicative of an inverted-U shape (p=0.005 for VTA-DA, p=0.008 for NB-ChAT, p=0.45 for SNc-DA for second degree polynomial of delay period fluorescence quintile; n=10 recording sites for VTA-DA, n=13 for SNc-DA, n=18 for NB-ChAT). Additionally, the Sasabuchi-Lind-Mehlum tests for inverted-U further validated the inverted-U relationship between accuracy and delay period fluorescence in VTA-DA and NB-ChAT populations (for detail on tests, see Methods, Inverted-U quantification; see Supplementary Figure 10g). In contrast to the delay period, fluorescence was not related to accuracy with an inverted-U according to the same sets of tests in any of these regions during the sample and choice periods (see Supplementary Figure 10g for p-values of all tests of the inverted-U).

To control for the possibility that the rat’s position during the delay period could contribute to the inverted-U relationship between fluorescence and accuracy, we repeated the same analysis using the subset of the delay period data during which the animal’s head was near the nosepoke (Supplementary Figure 11). The significant inverted-U relationship between the delay period GCaMP fluorescence and accuracy in VTA-DA and NB-ChAT populations was maintained in this subset of the data (mixed-effect linear regression in which accuracy was predicted based on the first and second degree polynomial of delay period fluorescence quintile, delay period duration, and a random effect of individual recording site; p=0.003 for VTA-DA, p=0.09 for SNc-DA, p=0.002 NB-ChAT).

Thus, we observed a neural correlate of the inverted-U relationship between neuromodulation and short-term memory performance, specifically during the delay period. To our knowledge, a neural correlate of this phenomenon has not previously been reported.

### Optogenetic inhibition of VTA-DA neurons selectively impairs short-term memory, while inhibition of SNc-DA, NB-ChAT and MS-ChAT neurons does not

To determine if the activity we measured in neuromodulatory populations contributes causally to task performance, we optogenetically inhibited each population throughout a trial, on a subset of trials. To this end, we injected Cre-dependent NpHR into the VTA or SNc of TH::Cre rats and implanted bilateral fibers above the injection site (Figure 5a-b, Supplementary Figure 12-13).

**Figure 5.**
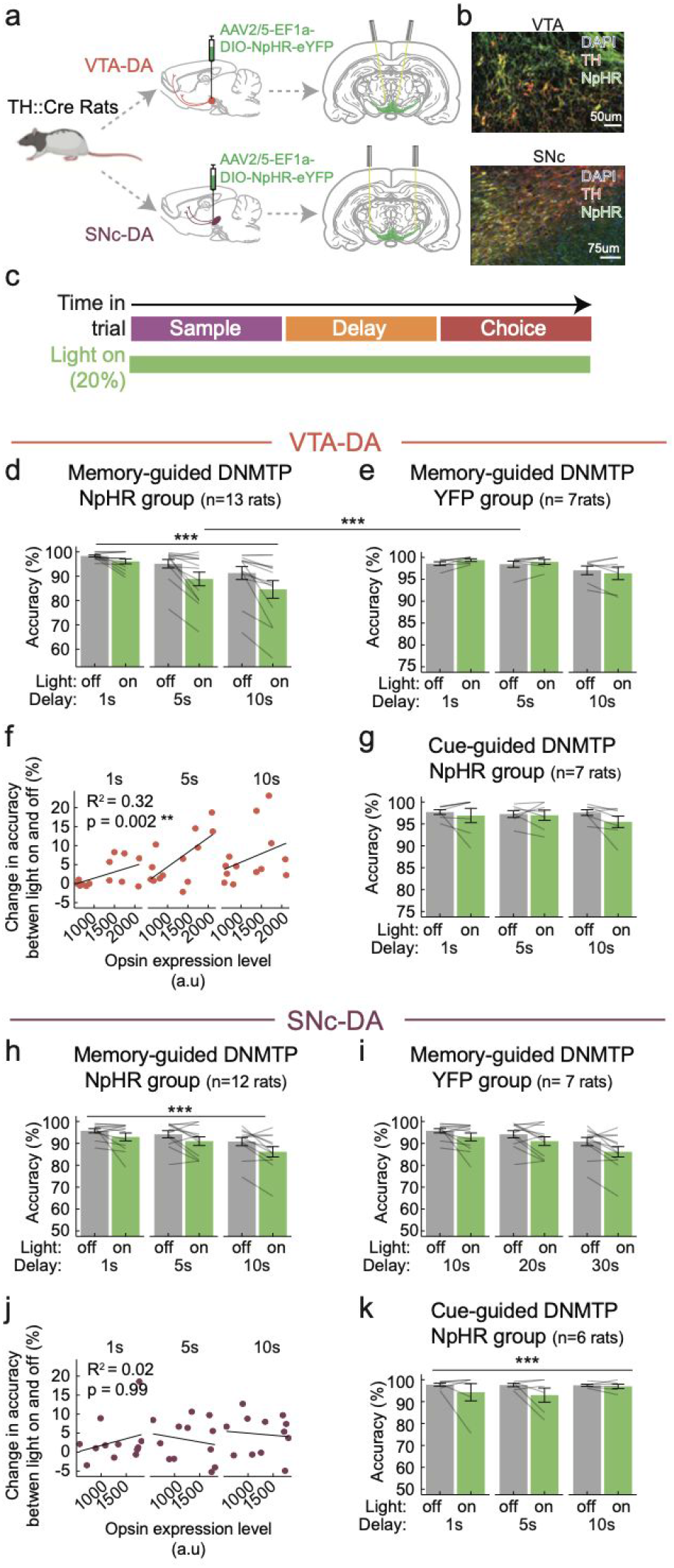
Optogenetic inhibition of VTA-DA, but not SNc-DA population, selectively impairs short-term memory. (a) Schematic of VTA-DA and SNc-DA targeting strategy using TH::Cre rats and Cre-dependent AAV2/5 DIO-NpHR-YFP (or DIO-YFP for the control group) virus injected into the VTA (top) or SNc (bottom), respectively. (b) Example histology from the VTA (top) and SNc (bottom), showing the co-localization of TH (red) and NpHR (green). (c) Schematic of experimental design for the entire-trial inhibition experiment. Continuous illumination (532nm at 5-6mW power at the fiber tip) was presented throughout the entirety of a trial, on a randomly selected 20% of trials, every other day. (d) Inhibition of VTA-DA neurons during the memory-guided DNMTP task impaired accuracy (mixed-effect logistic regression, correct/incorrect choice predicted based on fixed effects of light, delay duration, and random effect of individual rat; p<0.001 for light, n=13 NpHR rats). (e) In YFP control animals, the effect of light was not significant (mixed-effect logistic regression, correct/incorrect choice predicted based on fixed effects of light, delay duration, and random effect of individual rat; p=0.23 for light; n=7 YFP rats) and there was a significant interaction between laser x group (left and right combined: mixed-effect logistic regression, correct/incorrect choice predicted based on fixed effects of group, light, delay duration, and random effect of individual rat; p<0.001 for light x group, n=13 NpHR + 7 YFP rats). (f) In the VTA-DA NpHR group, virus expression level as measured by fluorescence intensity correlates with the behavioral effect size as measured by change in accuracy between light-on and light-off trials for each rat (y-axis, two-way ANOVA, light impairment in accuracy explained by opsin expression level and delay duration; p=0.002 for opsin expression, p=0.07 for delay duration; R2=0.32; n=13 NpHR rats). (g) Optogenetic inhibition of VTA-DA neurons during the cue-guided DNMTP task. Accuracy (y-axis) was not impaired in light-on trials (green bar) compared to light-off trials (gray bar; mixed-effect logistic regression, correct/incorrect choice predicted based on fixed effects of light, delay duration, and random effect of individual rat; p=0.14 for light; n=7 NpHR rats). The accuracy impairment in the memory-guided but not cue-guided DNMTP task suggests that the effect of VTA-DA inactivation was specifically attributable to the short-term memory component of the task. (h-k) same as (d-g) but in the SNc-DA group. Unlike VTA-DA, the accuracy was impaired in both the memory-guided (mixed-effect logistic regression, correct/incorrect choice predicted based on fixed effects of light, delay duration, and random effect of individual rat; p<0.001 for light; n=12 NpHR rats) and cue-guided DNMTP task (mixed-effect logistic regression, correct/incorrect choice predicted based on fixed effects of light, delay duration, and random effect of individual rat; p<0.001 for light; n=6 NpHR rats), suggesting that the effect of SNc-DA inactivation cannot be specifically attributable to the short-term memory.

Full-trial inhibition of DA neurons in the VTA significantly impaired rats’ performance in the memory-guided DNMTP task (Figure 5c-d, mixed-effect logistic regression, correct/incorrect choice predicted based on fixed effects of light, delay, and random effect of individual rat; p < 0.001 for light; n=13 NpHR rats). The level of opsin expression (as assessed by fluorescence intensity in histology) was significantly correlated with the optogenetic impairment (Figure 5f, two-way ANOVA light impairment explained by fluorescence level and delay; p=0.002 for fluorescence, p=0.07 for delay; R^2^=0.32; n=13 NpHR rats). There was no significant light-induced impairment in the YFP control animals (Figure 5e, mixed-effect logistic regression, correct/incorrect choice predicted based on fixed effects of light, delay and random effect of individual rat; p=0.23 for light; n=7 YFP rats, Supplementary Figure 12b), and there was a significant light x group interaction between the NpHR and YFP groups (Figure 5d-e, mixed-effect logistic regression, correct/incorrect choice predicted based on fixed effects of light, delay, group, and random effect of individual rat; p<0.001 for light x group; n=13 NpHR, 7 YFP rats). In contrast to the effect on accuracy, choice omission rate did not show significant light-induced change (Supplementary Figure 14).

To determine if the impairment induced by VTA-DA inhibition was specifically attributable to the short-term memory component of the task, we compared performance to a control variant of the task, in which rats did not have to use short-term memory (Figure 5g). In the cue-guided task, the motor requirements were identical, but a light cue directly above the correct choice lever was illuminated during the choice period, to signal which lever was correct. Optogenetic inhibition of DA neurons in VTA did not affect performance in the cue-guided task (Figure 5g, mixed-effect logistic regression, correct/incorrect choice predicted based on fixed effects of light, delay and random effect of individual rat; p=0.14 for light; n=7 NpHR rats, Supplementary Figure 12c). Thus, the effect of optogenetic inhibition of VTA-DA appeared to be dependent on the task having a short-term memory component.

Next, we investigated if SNc-DA neurons also contributed causally to short-term memory. We performed an identical set of inhibition experiments in the SNc as we had in the VTA. Optogenetic inhibition of SNc-DA neurons impaired accuracy in the memory-guided task (Figure 5h, mixed-effect logistic regression, correct/incorrect choice predicted based on fixed effects of light, delay and random effect of individual rat; p<0.001 for light; n=12 NpHR rats, Supplementary Figure 13a). This effect was not present in control rats expressing YFP in SNc-DA (Figure 5i, mixed-effect logistic regression, correct/incorrect choice predicted based on fixed effects of light, delay and random effect of individual rat; p = 0.12 for light; n = 7 YFP rats, Supplementary Figure 13b). However, unlike VTA-DA, there was no significant interaction between light on/off and opsin/yfp group (Figure 5h-i, mixed-effect logistic regression, correct/incorrect choice predicted based on fixed effects of group, light, delay, and random effect of individual rat; p =0.27 for light x group; n=12 NpHR, 7 YFP rats), and the level of opsin expression did not correlate with the optogenetic impairment (Figure 5j, two-way ANOVA, light impairment explained by fluorescence level and delay; p=0.99 for fluorescence, p=0.69 for delay; R^2^=0.02; n=12 NpHR rats).

Moreover, unlike VTA-DA neurons, optogenetic inhibition of SNc-DA neurons during the control cue-guided task significantly impaired accuracy (Figure 5k, mixed-effect logistic regression, correct/incorrect choice predicted based on fixed effects of light, delay and random effect of individual rat; p < 0.001 for light; n = 6 NpHR rats, Supplementary Figure 13c). The presence of light effect in *both* the memory-guided and cue-guided tasks suggests that the effect of SNc-DA inhibition is different from that of VTA-DA and cannot be specifically attributed as a short-term memory deficit.

We next asked if ascending ChAT neurons in the NB and MS contributed causally to short-term memory (Figure 6). To address this, throughout the trial on a subset of trials, we inhibited ChAT neurons in the MS and NB in ChAT::Cre rats performing the DNMTP task (Figure 6a-b; Supplementary Figure 15-16). We found that the inhibition of neither ChAT population affected short-term memory performance (Figure 6c, NB-ChAT group, mixed-effect logistic regression, correct/incorrect choice predicted based on fixed effects of light, delay and random effect of individual rat; p=0.9 for light; n=5 NpHR rats; Figure 6d, MS-ChAT group, mixed-effect logistic regression, correct/incorrect choice predicted based on fixed effects of light, delay and random effect of individual rat; p=0.17 for light; n=3 NpHR rats).

**Figure 6.**
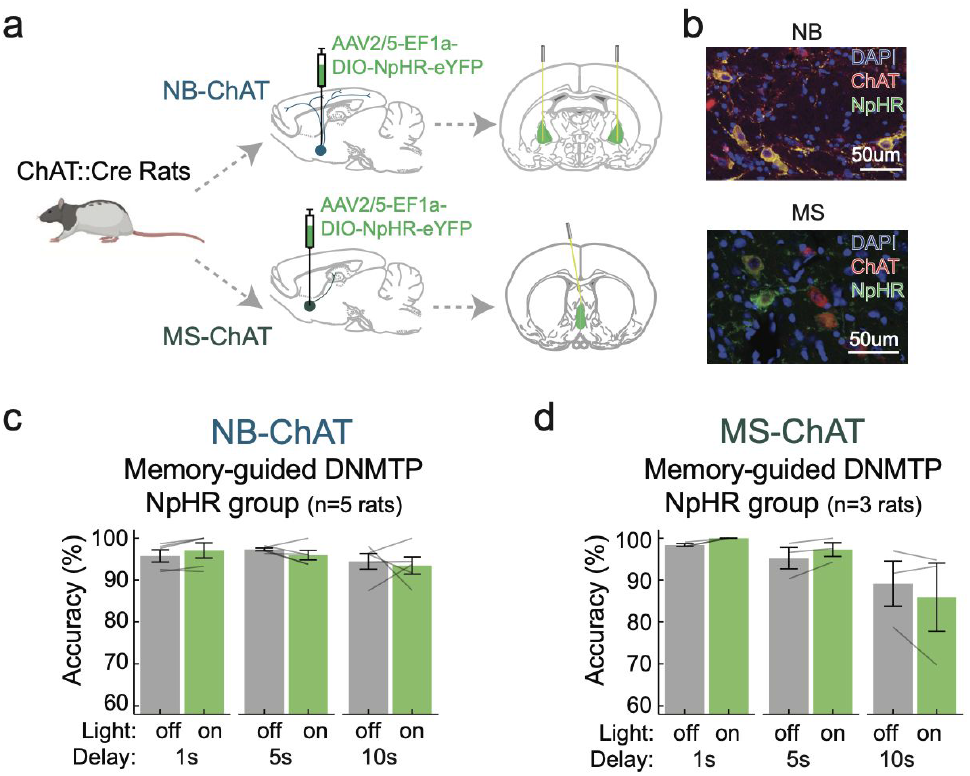
Optogenetic inhibition of NB-ChAT and MS-ChAT does not impair short-term memory. (a) Schematic of NB-ChAT and MS-ChAT targeting strategy using ChAT::Cre rats and AAV2/5 DIO-NpHR-YFP virus injected into the NB (top) or MS (bottom). (b) Example histology from the NB (top) and MS (bottom), showing co-localization of ChAT (red) and NpHR (green) (c) Optogenetic inhibition of NB-ChAT does not affect performance in the DNMTP task (mixed-effect logistic regression, correct/incorrect choice predicted based on fixed effects of light, delay duration and random effect of individual rat, p=0.9 for light; n=5 NpHR rats). (d) Optogenetic inhibition of MS-ChAT does not affect short-term memory (mixed-effect logistic regression, correct/incorrect choice predicted based on fixed effects of light, delay duration and random effect of individual rat; p=0.17 for light; n=3 NpHR rats).

Taken together, this suggests that the causal contribution of VTA-DA to short-term memory is unique relative to the other neuromodulatory populations that we examined.

### Optogenetic inhibition of VTA-DA neurons during the delay period most severely impairs short-term memory

After determining that VTA-DA neurons contribute to short-term memory, we next asked *when* they do so - during the sample, delay, or choice period of the task? To address this, on a subset of trials, and in a randomly interleaved manner, we inhibited VTA-DA neurons during one of the three epochs (Figure 7a; Supplementary Figure 12a).

**Figure 7.**
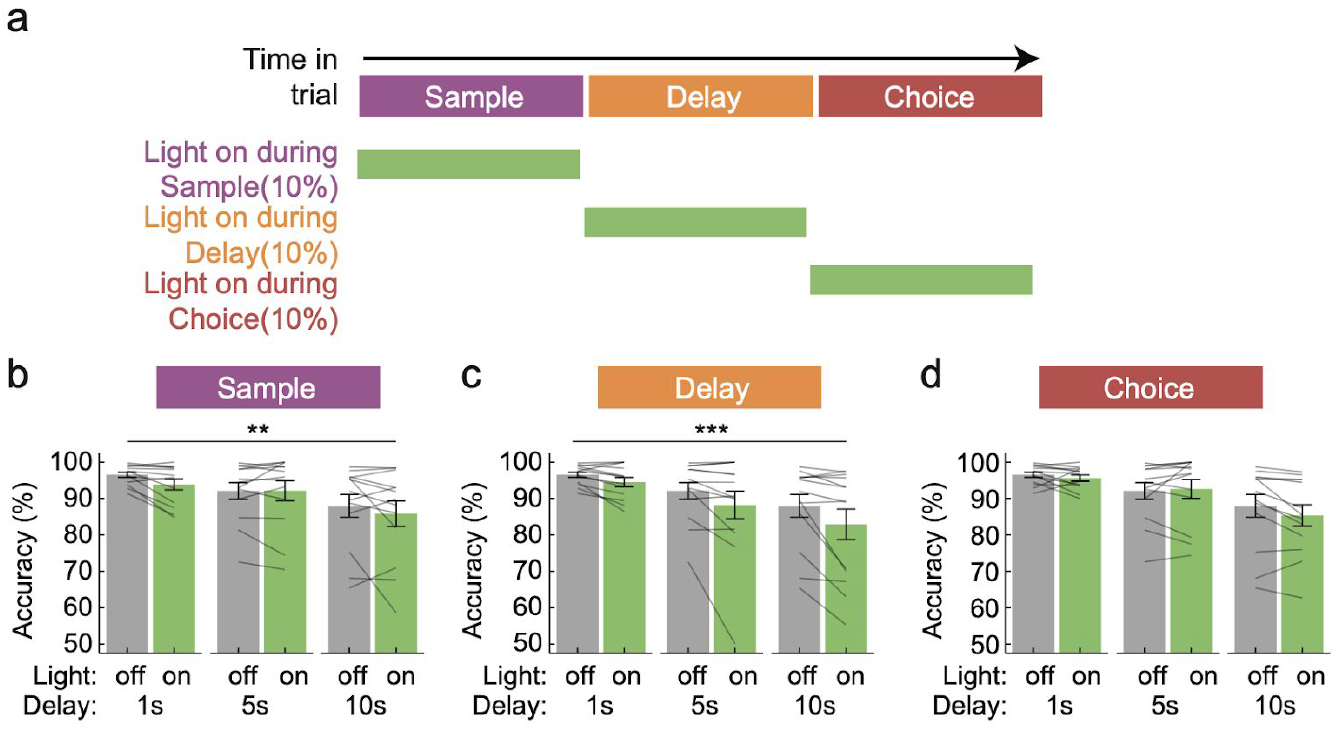
Optogenetic inhibition of VTA-DA during the sample or delay, but not choice, produces impairments in short-term memory. (a) Schematic of experimental design for sub-trial inhibition. Continuous light was presented during either the sample, delay or choice periods of a trial in an interleaved manner, with each manipulation occurring on 10% of trials (532 nm, ~5-6mW at the fiber tip). (b-d) Accuracy for light-on (green bar) vs light-off (gray bar) trials for the sample (b), delay (c), and choice (d) period. Mixed-effect logistic regression (n=13 NpHR rats) to predict correct/incorrect choice based on fixed effects of light (light off, light during sample, light during delay, light during choice), delay duration, and random effect of individual rat reveals a significant effect of light during sample (p= 0.002) and delay (p<0.001) but not choice (p=0.3).

Optogenetic inhibition of VTA-DA neurons during the sample and delay period, but not the choice period, significantly impaired short-term memory (Figure 7b-d, mixed-effect logistic regression, correct/incorrect choice predicted based on fixed effects of light epoch, delay, and random effect of individual rat; p=0.002 for light on sample, p<0.001 for light on delay, p=0.3 for light on choice; n=13 NpHR rats). Note that the regression coefficient corresponding to light during the delay period was larger than that for the sample period, suggesting a bigger effect size for inhibiting during the delay than the sample period (sample: *β*=−0.25+0.081, delay: *β*=−0.45+0.076).

### Optogenetic activation of VTA-DA neurons during the delay, but not the sample period, impairs short-term memory

The inverted-U hypothesis posits that too much or too little DA would impair short-term memory. This would suggest that not only inhibition, but also activation of VTA-DA neurons would impair short-term memory. On the other hand, the gating theory would suggest that more DA during the sample period could enhance short-term memory. To our knowledge, these ideas have not been tested with direct manipulation of DA neural activity at sub-trial resolution.

Thus, we next injected an AAV2/5 expressing Cre-dependent ChR2 into the VTA of TH::Cre rats (Figure 8a-b, Supplementary Figure 12d). We briefly activated VTA-DA neurons at the time of the sample presentation, to simulate the phasic response observed with fiber photometry (Figure 8c; 5ms pulse duration, 5 pulses/s of 20 Hz stimulation, ~15mW). Optogenetic activation did not improve short-term memory, which was *not* consistent with predictions from the gating theory (Figure 8d, mixed-effect logistic regression, correct/incorrect choice predicted based on fixed effects of light, delay and random effect of individual rat; p= 0.13 for light; n=9 ChR2 rats; Supplementary Figure 10b).

**Figure 8.**
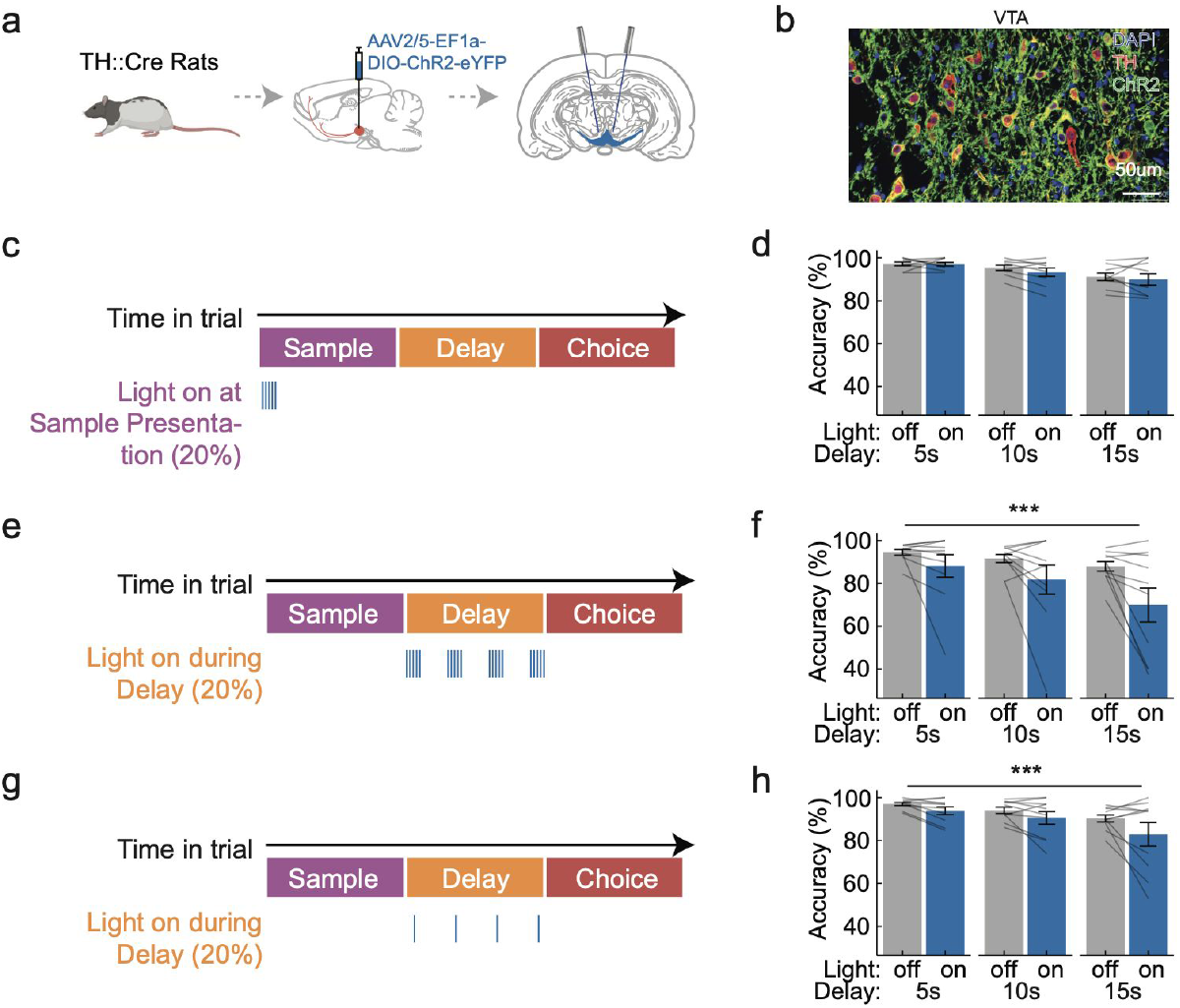
Optogenetic activation of VTA DA neurons during the delay, but not sample, impairs short-term memory. (a) Schematic of VTA-DA targeting strategy using TH::Cre rats and AAV2/5 DIO-ChR2-YFP virus. (b) Histology of ChR2 expression in VTA-DA neurons. (c) Schematic of experimental design for optogenetic activation at sample lever presentation. VTA-DA was activated using a burst of 5 pulses at 20Hz at the sample lever presentation 447nm, 5ms pulse duration, ~15mW light power) (d) Performance in DNMTP task for light-on (blue bar) vs light-off (gray bar) trials for VTA-DA activation using the protocol described in (c). VTA-DA activation during sample presentation did not modulate accuracy (mixed-effect logistic regression, correct/incorrect choice predicted based on fixed effects of light, delay duration, and random effect of individual rat; p=0.13 for light; n=9 ChR2 rats). (e) Schematic of experimental design for burst activation of VTA-DA during the delay period (5ms pulse duration, 5 pulses at 20Hz per burst, 1 burst/s). (f) Performance in the DNMTP task for light-on (blue bar) vs light-off (gray bar) trials for VTA-DA activation during the delay period using the protocol described in (e). VTA-DA activation in bursts during the delay period significantly impaired accuracy. (mixed-effect logistic regression, correct/incorrect choice predicted based on fixed effects of light, delay duration, and random effect of individual rat; p<0.001 for light; n=10 ChR2 rats) (g) Schematic of experimental design for tonic optogenetic activation of VTA-DA during the delay period (5ms pulse duration, 1 pulse/s). (h) Performance in the DNMTP task for light-on (blue bar) vs light-off (gray bar) trials for VTA-DA activation during the delay period using the protocol described in (g). VTA-DA tonic activation during the delay period significantly impaired accuracy. (mixed-effect logistic regression, correct/incorrect choice predicted based on fixed effects of light, delay and random effect of individual rat; p<0.001 for light; n=10 ChR2 rats).

Next, we activated VTA-DA neurons during the delay period, which is when we observed the most severe impairment of short-term memory from the optogenetic inhibition (Figure 8e). We found that the optogenetic activation of VTA-DA neurons resulted in significant impairment of task performance, similar to our results from inhibition of this population (Figure 8f, mixed-effect logistic regression, correct/incorrect choice predicted based on fixed effects of light, delay and random effect of individual rat; p<0.001 for light; n=10 ChR2 rats; Supplementary Figure 12d). In fact, even mild optogenetic activation (1 pulse/s, 5ms pulse duration) of VTA-DA neurons during the delay period resulted in a significant impairment in performance (Figure 8g-h, mixed-effect logistic regression, correct/incorrect choice predicted based on fixed effects of light, delay and random effect of individual rat; p<0.001 for light; n=10 ChR2 rats).

Taken together, we conclude that activation or inhibition of VTA-DA neurons during the delay period impairs short-term memory performance, despite the depressed activity in this population during that time.

## Discussion

### New support for an inverted-U relationship between DA and short-term memory maintenance

Midbrain DA neurons are known to respond to reward-predicting cues and unexpected rewards - in other words, they encode errors in the prediction of reward (Bayer and Glimcher, 2005; Cohen et al., 2012; Roesch et al., 2007; Schultz, 1986, 1998; Schultz et al., 1997). In addition, DA has been implicated in short-term memory, primarily through pharmacological manipulations in monkeys (Arnsten et al., 1994; Cai and Arnsten, 1997; Sawaguchi and Goldman-Rakic, 1991; Williams and Goldman-Rakic, 1995). However, it has been unclear if and how to integrate these literatures. In particular, pharmacological experiments had suggested that DA is most important to short-term memory during the delay period (Figure 1a; Vijayraghavan et al., 2007; Williams and Goldman-Rakic, 1995). Since there are usually no reward-predicting cues or rewards during the delay period, it is not obvious if and why there would be DA activity at that time.

Thus, to reconcile the role of DA in reinforcement learning with one in short-term memory, it was proposed that DA contributes to the *updating* of short-term memory with new information (the ‘gating’ theory; Figure 1b; Braver and Cohen, 1999, 2000; O’Reilly and Frank, 2006), which should occur at the time of reward-predicting stimuli, rather than to the *maintenance* of short-term memory during the delay period, which was the original hypothesis from pharmacological experiments. Since pharmacology is too slow to distinguish between a role in updating versus maintaining short-term memory, these hypotheses have remained untested. Thus, a major goal of this study was to directly measure and manipulate DA neuron activity during a short-term memory task with a distinct “sample period” in which short-term memory is updated, as well as a “delay period” in which short-term memory is maintained, to determine which aspect of short-term memory DA supports.

Our recordings revealed activity in DA neurons that was consistent with reward prediction error and therefore the ‘gating theory’. We observed relatively low activity in VTA-DA neurons during the delay period, and elevated activity during the sample, choice and outcome period. This is consistent with VTA-DA responses primarily being explained by reward prediction error: reward-predicting cues appear during the sample and choice period (the lever presentation), and reward occurs during the outcome period. In addition, the choice lever presentation elicited higher activity than the sample lever presentation, which is also consistent with a reward prediction error, assuming a temporally discounted reward expectation function (Fiorillo et al., 2008; Kobayashi and Schultz, 2008; Mazur, 1987; Richards et al., 1997; Roesch et al., 2007; Samuelson, 1937; Starkweather et al., 2017). Similar to VTA-DA, SNc-DA neurons also did not have elevated activity during the delay period, although the activity was not as low as VTA-DA.

Based on these neural correlates, we expected that DA might be causally involved in the sample period of short-term memory, consistent with the gating theory. In fact, we did observe a mild impairment in short-term memory as a result of inhibiting during the sample period, providing some causal support for that hypothesis. However, this effect was relatively small, and we observed no effect of activation during this period.

In addition to providing some new support for the gating theory, our recordings also revealed new insights about the relationship between endogenous DA activity and short-term memory performance. To our surprise, we observed an “inverted-U” relationship between the delay period activity in VTA-DA and cognitive performance. To our knowledge, this is the first evidence that the activity of any neuromodulator relates to performance with an inverted-U. This correlational evidence provides a new form of support of classic ideas that had emerged from pharmacological manipulations, which had artificially manipulated receptor activation but provided no insight into the natural activity patterns. This association between accuracy and dopamine may relate to roles that have been ascribed for dopamine in regulating motivation or internal state (Berridge and Robinson, 1998; Bromberg-Martin et al., 2010; Cohn et al., 2015; Hamid et al., 2016; Lammel et al., 2012; Matsumoto and Hikosaka, 2009; Tye et al., 2013; Vander Weele et al., 2018; Westbrook and Braver, 2016).

In addition, bi-directional optogenetic manipulations revealed that the delay period was most relevant to short-term memory, as inhibition or activation led to relatively large impairments in performance, despite the low activity at that time. Thus, our manipulation of cell bodies very much resembled the dose-dependent “inverted-U” effects of D1 agonist treatment in monkey PFC during spatial short-term memory (Cai and Arnsten, 1997; Murphy et al., 1996; Sawaguchi and Goldman-Rakic, 1991; Vijayraghavan et al., 2007; Williams and Goldman-Rakic, 1995; Zahrt et al., 1997). These findings highlight a dissociation between when DA neurons are most active and when their activity most affects short-term memory, and reveal a new form of correlational and causal support that are consistent with classic ideas of an “inverted-U” relationship between DA and cognition. Given that the reward prediction error framework does not predict modulation in DA activity during the delay period, these results suggest that that framework is insufficient to fully explain DA function in short-term memory.

### Previous work measuring DA neural activity or DA efflux during short-term memory

A previous paper by (Phillips et al., 2004) measured DA efflux with microdialysis during a memory task with considerably longer delays (30min, 1hr, 6hr). The major finding of that study was a negative correlation between delay duration and DA release during the choice period, which we also report here (Supplementary Figure 5).

(Matsumoto and Takada, 2013) recorded from VTA-DA and SNc-DA in non-human primates in a short-term memory task, and found that SNc-DA responded to the sample stimulus only when the subject was required to store it in short-term memory. From these neural correlates, they concluded that only SNc-DA (but not VTA-DA) activity reflects short-term memory demand. However, they did not manipulate neural activity in these populations to assess causality, and in fact our finding that VTA-DA and not SNc-DA contribute to WM, and do so preferentially during the delay period, provides another potential interpretation of their results. Specifically, our results suggest that the SNc-DA response to the sample stimulus observed in their study may only be correlational and not causal to short-term memory.

Although we compared effects of VTA-DA and SNc-DA inhibition in a short-term memory task and a cue-guided task (Figure 5), we did not directly compare neural correlates in the DA system of these two tasks. Interestingly, (Watanabe et al., 1997) used in vivo microdialysis to demonstrate an increase in DA level in the principal sulcus in primates after performance of a short-term memory task but not a cue-guided task. Whether such differences exist in the fast dynamics of VTA-DA activity within a trial remains to be established.

The task we used in this paper is conceptually similar to the delayed response task used in the original Goldman-Rakic papers (Arnsten et al., 1994; Sawaguchi and Goldman-Rakic, 1991; Williams and Goldman-Rakic, 1995) that implicated DA in short-term memory. In both our task and the task used in those original papers, there is a stimulus that is updated into short-term memory during a sample period, then a delay period when it is retained, and finally a choice period that serves as a readout of the memory. Thus, we could leverage this task to examine the contribution of each neuromodulatory population to each epoch of short-term memory. Of note, there are other more powerful working memory tasks that have been applied in humans and non-human primates, such as the N-back task and the AX-CPT task (Blackman et al., 2016; D’Ardenne et al., 2012; Kirchner, 1958; Owen et al., 2005; Rosvold et al., 1956). These tasks allow better differentiation between retrospective and prospective memory, as well as classification of correct and incorrect trials into two types (hit vs correct rejection, false alarm vs miss). In addition, the N-back task allows an examination of the effect of working memory load on performance.

### Distinctions and similarities in neural correlates of short-term memory across DA and ChAT populations

Aside from clarifying the temporal contribution of DA to short-term memory, another major goal of this work was to directly compare the dopaminergic and cholinergic contribution to short-term memory. Only a few studies have characterized basal forebrain cholinergic neurons during behavior (Hangya et al., 2015; Harrison et al., 2016), and therefore the similarity of activity of dopaminergic and cholinergic populations remains underexplored not only in short-term memory tasks, but also more generally.

Perhaps the most prominent difference between the populations we examined was the MS-ChAT neurons, which encoded speed much more than the other populations. To our knowledge, preferential encoding of speed in this population has not previously been reported. Additionally, we found that VTA-DA preferentially responded to lever presentation cue whereas NB-ChAT preferentially responded to lever press action during the sample and choice periods.

The most striking similarity we observed was across the three task-encoding populations (NB-ChAT, VTA-DA, SNc-DA), all of which had reward responses and elevated activity during the sample and choice periods. This is consistent with previous reports of reward responses not only in DA neurons but also NB-ChAT neurons (Hangya et al., 2015; Teles-Grilo Ruivo et al., 2017). Another similarity was between the NB-ChAT and VTA-DA population, which both had an inverted-U shaped relationship between delay period activity and performance.

However, note that since we did not record from single neurons, our conclusions pertain to the average signal in each region. Accordingly, we cannot rule out the possibility that results may be dominated by a subset of strongly responding neurons.

### SNc-DA, NB and MS ChAT neurons do not contribute causally and selectively to short-term memory

In contrast to some of the similarities we observed in the neural correlates of the task across the neuromodulatory populations we examined, the causal contributions were more distinct. Only VTA-DA neurons contributed selectively to the short-term memory task, as SNc-DA inhibition affected both the short-term memory task and a control task. In addition, NB-ChAT and MS-ChAT populations were not causally involved in short-term memory.

The lack of involvement of the NB-ChAT populations is not aligned with classic lesion studies that used non-specific excitotoxins to lesion NB (i.e. ibotenic acid, quisqualic acid) and reported deficits in a battery of spatial memory tests such as the Morris water maze (Connor et al., 1991; Mandel and Thal, 1988; Mandel et al., 1989), and radial maze (Hodges et al., 1989; Lerer and Warner, 1986; Turner et al., 1992). However, our negative result with NB-ChAT population is consistent with subsequent and more specific studies with cholinergic neuron-selective neurotoxin, IgG-saporin (Baxter and Bucci, 2013; Baxter et al., 1995; Torres et al., 1994; Wenk et al., 1994).

In contrast to NB-ChAT neurons, MS-ChAT neurons have been implicated in certain spatial short-term memory tasks with IgG-saporin (Torres et al., 1994). However, our neural correlate demonstrated that MS-ChAT population primarily encodes the animal’s movement rather than task events, providing little reason to believe that these neurons would be selectively involved in short-term memory. One possibility may be that the septo-hippocampal ChAT pathway is only selectively involved in short-term memory in the case of novel stimuli (Hasselmo and Sarter, 2011; Hasselmo and Stern, 2006), perhaps by contributing to the generation of exploratory behavior.

## Supplementary Figures

**Supplementary Figure 1.**
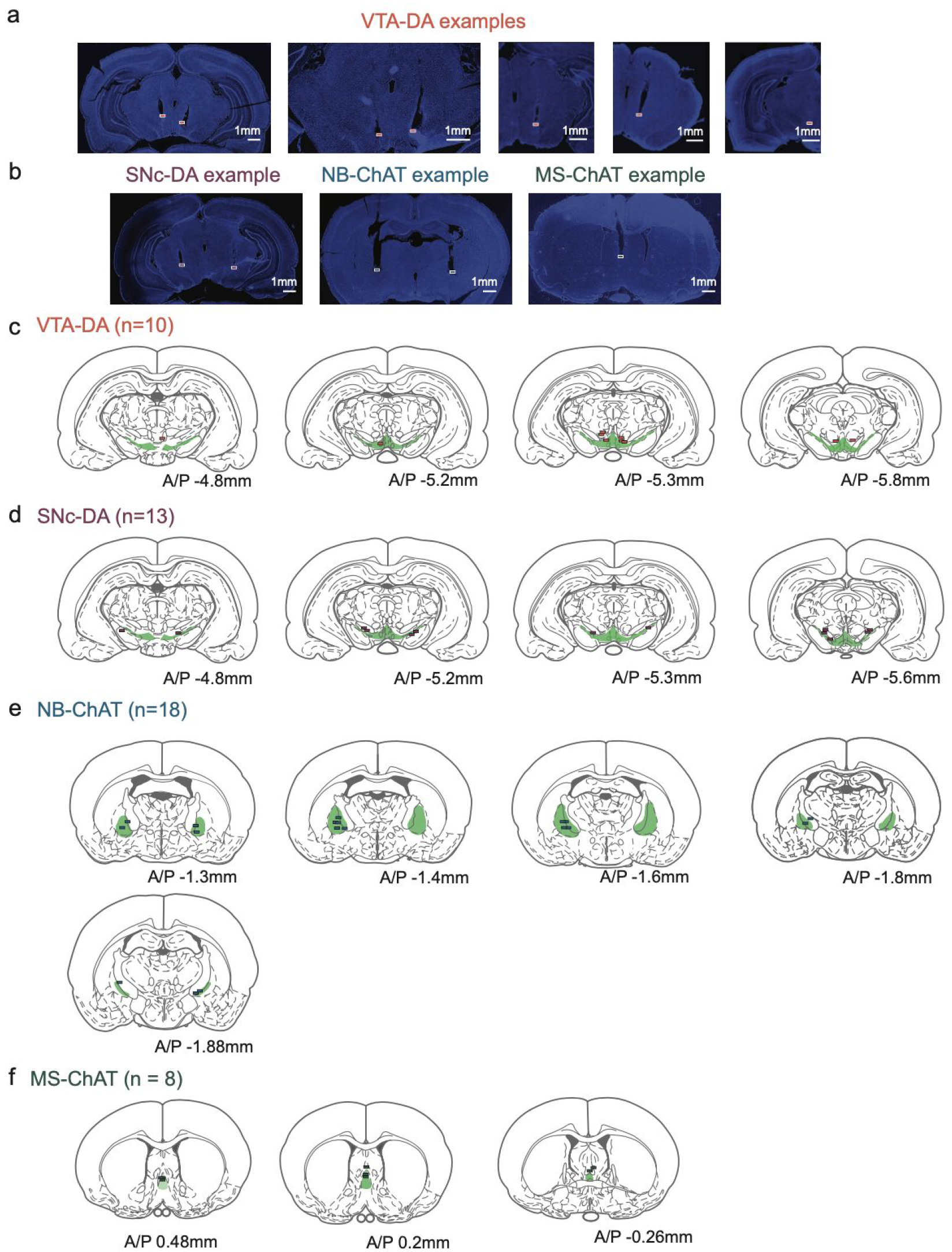
Summary of fiber photometry recording sites. (a) Examples of VTA-DA fiber placements (n=4 rats). (b) Example histology images showing the fiber placement from SNc-DA, NB-ChAT and MS-ChAT recordings for data in Figures 2-4. (c) VTA-DA fiber photometry recording sites (n=10 recording sites). Each line represents the reconstructed location of a fiber tip from histology. Green shaded area is VTA/SNc. (d) same as (c) but from SNc-DA fiber photometry recording sites (n=13 recording sites). (e) same as (c) but from NB-ChAT fiber photometry recording sites (n=18 recording sites). Green shaded area is NB. (f) same as (c) but from MS-ChAT fiber photometry recording sites (n=8 recording sites). Green shaded area is MS.

**Supplementary Figure 2.**
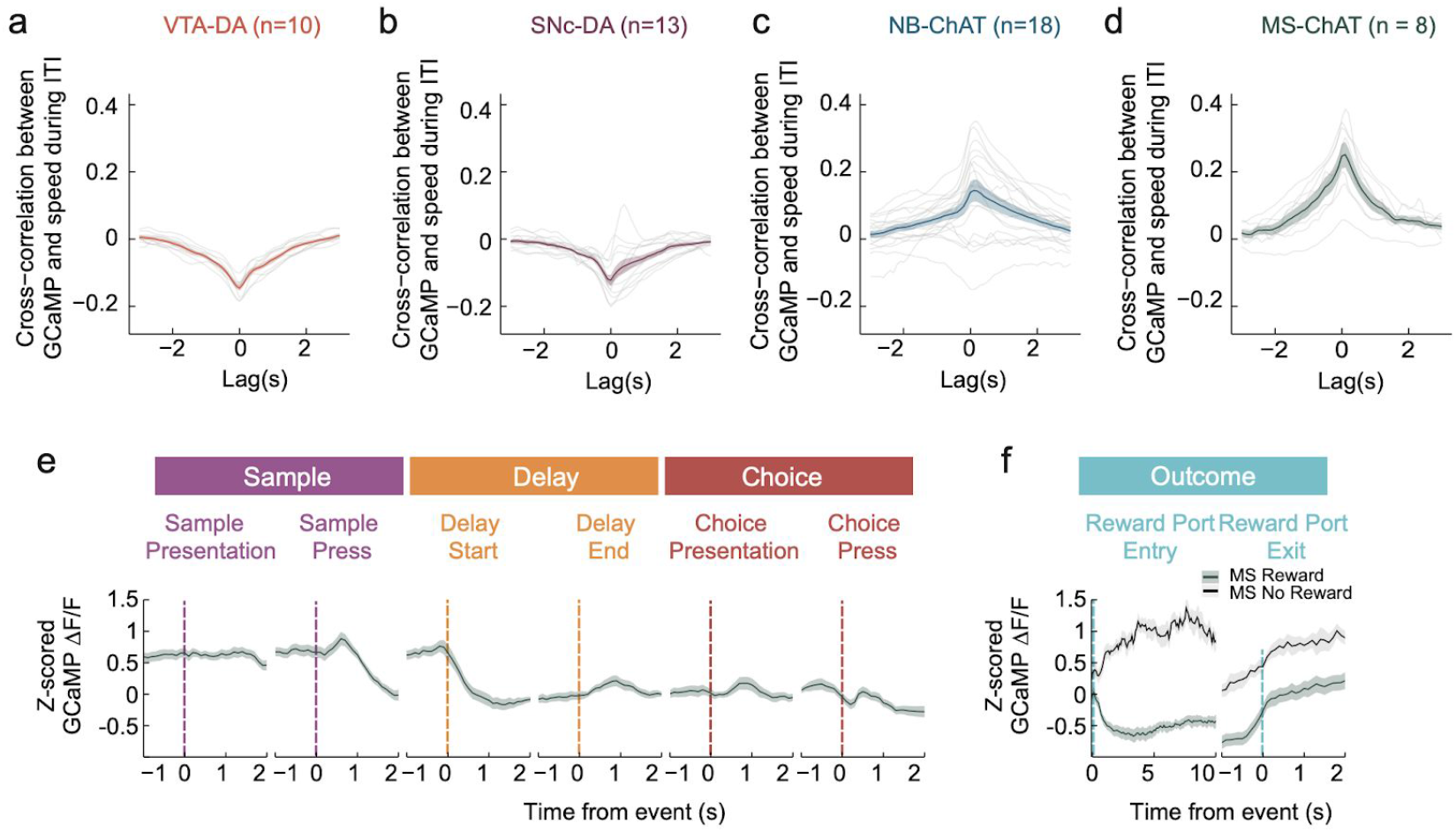
During ITI, GCaMP fluorescence is best correlated with speed in MS-ChAT compared to VTA-DA, SNc-DA, and NB-ChAT populations. Cross-correlation between GCaMP fluorescence and speed during the inter-trial interval (ITI) in VTA-DA (a), SNc-DA (b), NB-ChAT(c), and MS-ChAT (d) populations (gray lines show individual recordings, colored line and shade are mean+sem across recordings, n=10 recording sites for VTA-DA, n=13 recording sites for SNc-DA, n=18 recording sites for NB-ChAT and n=8 recording sites for MS-ChAT). DA subpopulations show negative correlation between GCaMP fluorescence and speed during ITI. In contrast, ChAT subpopulations show positive correlation between GCaMP and speed during the ITI. Notably, MS-ChAT population shows the highest correlation between GCaMP and speed on average compared to the other three populations. (e) Z-scored GCaMP fluorescence from MS-ChAT recordings time-locked to each task event during the sample, delay and choice periods (mean+sem across recordings, n=8 recording sites). Data from all 10s delay trials. (f) Z-scored GCaMP fluorescence from MS-ChAT recordings during the outcome period, separated by rewarded and unrewarded trials (mean+sem across recording sites, n=8 recording sites). Most of the variance observed in event time-locked activity is depressed GCaMP fluorescence during the delay period and reward consumption in rewarded trials (but not in unrewarded trials). Since rats tend to be stationary during the delay period to consume reward on rewarded trials, the observed depressed GCaMP fluorescence is consistent with the positive correlation with speed in this population.

**Supplementary Figure 3.**
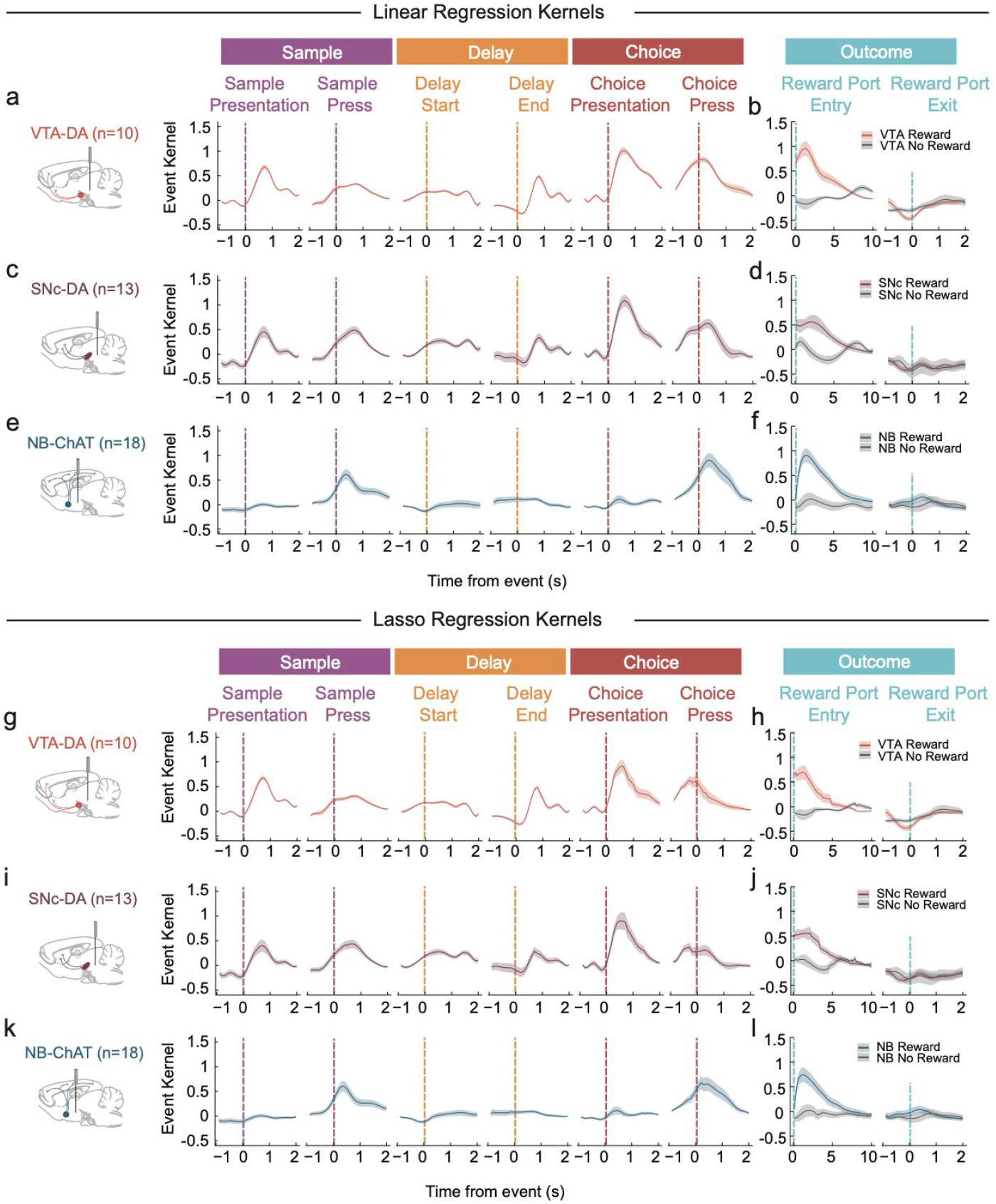
Response kernels for all task events in task event + speed model generated from linear regression and lasso (regularized) regression. (a-b) Response kernels (components of neural response that can be attributed to each task event based on the linear encoding model) is plotted for VTA-DA recordings (mean+sem across recordings, n=10 recording sites). In the encoding model, GCaMP was predicted by task events convolved with a spline basis set. (b). (c-d) same as (a-b) but for SNc-DA recordings (mean+sem across recordings, n=13 recording sites). (e-f) same as (a-b) but for NB-ChAT recordings (mean+sem across recordings, n=18 recording sites). (g-h) same as (a-b) but the response kernels are generated from using lasso regression, which regularizes predictors to prevent overfitting (n=10 recording sites). (i-j) same as (c-d) but using lasso regression in SNc-DA recordings (n=13 recording sites). (k-l) same as (e-f) but using lasso regression in NB-ChAT recordings (n=18 recording sites).

**Supplementary Figure 4.**
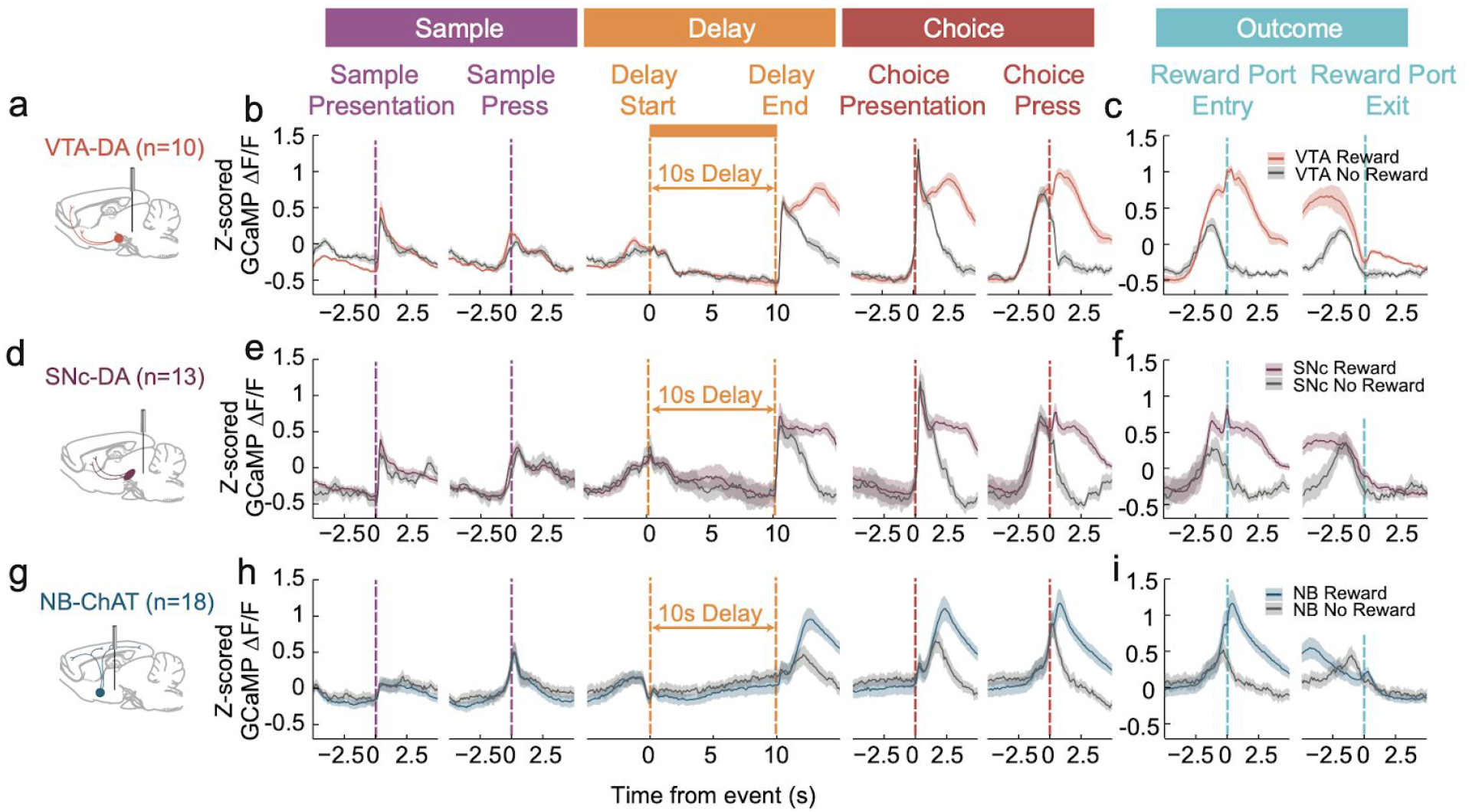
Difference in GCaMP fluorescence between rewarded versus unrewarded trials emerges after the presentation of reward cue at the time of choice press. Same data and plotting as Figure 4a-h, but here we separately plot rewarded and unrewarded trial data across all task events.

**Supplementary Figure 5.**
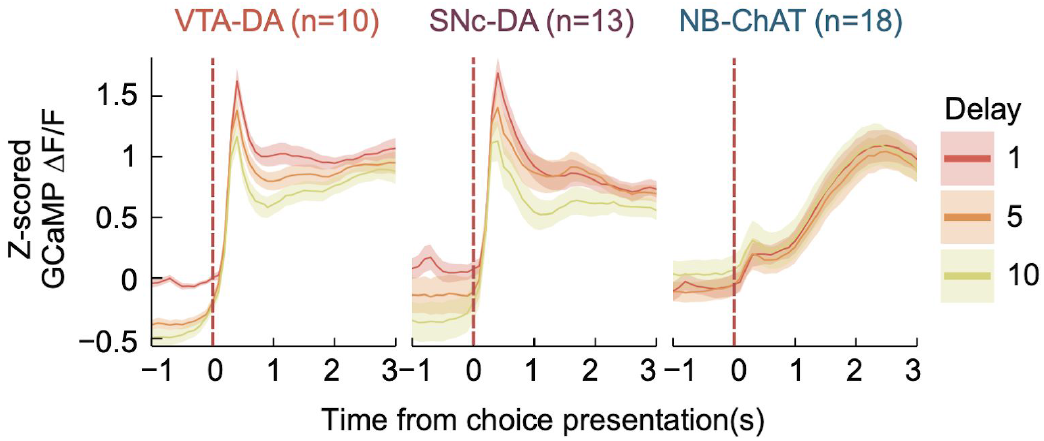
VTA-DA and SNc-DA choice period activity is negatively correlated with delay duration, consistent with reward prediction error. Z-scored GCaMP fluorescence time-locked to the beginning of choice period separated by delay duration for all trials in VTA-DA (left), SNc-DA (middle) and NB-ChAT populations. (mean+sem across recordings for each delay duration, n=10 recording sites for VTA-DA, n=13 recording sites for SNc-DA, n=18 recording sites for NB-ChAT).

**Supplementary Figure 6.**
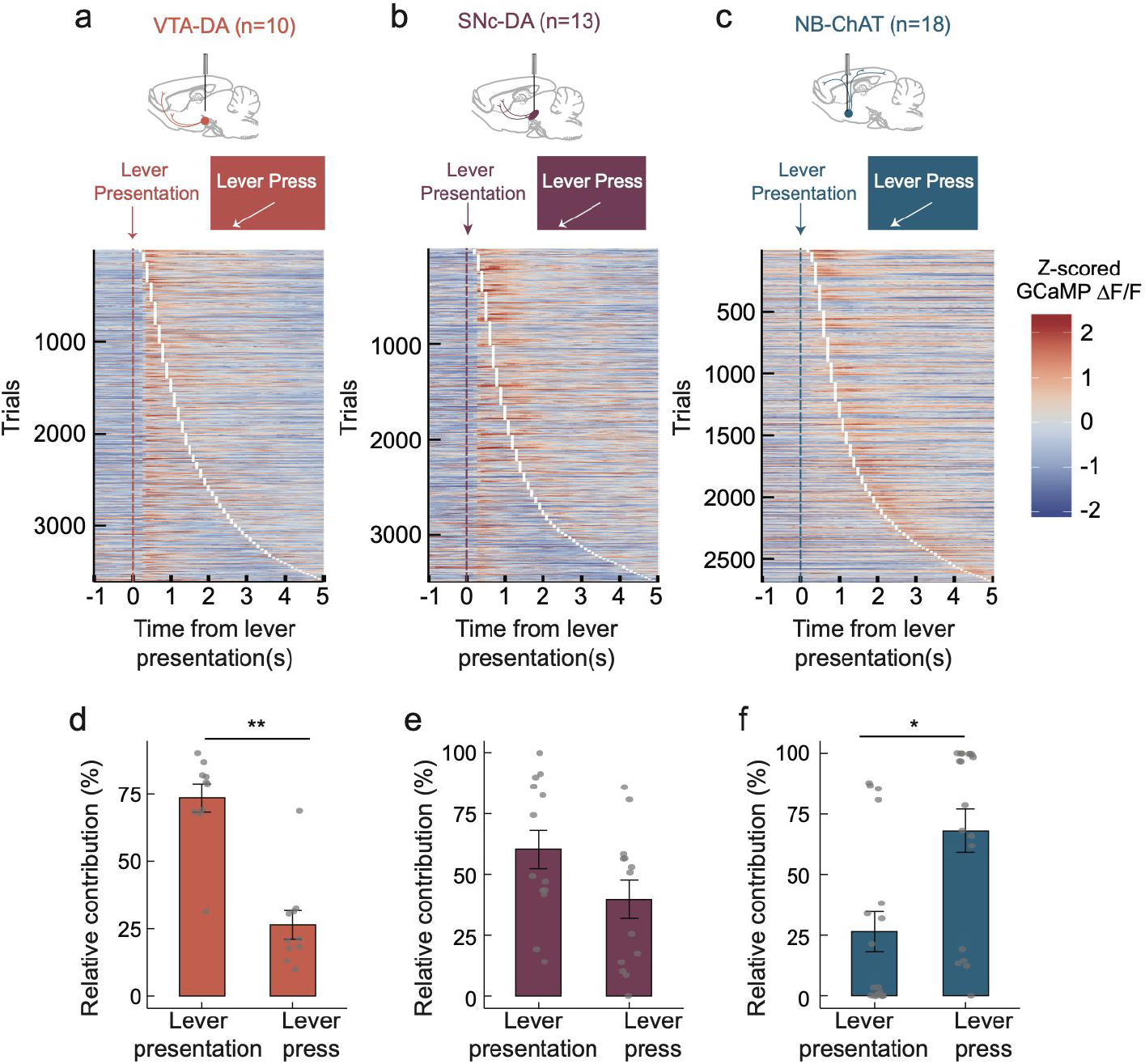
VTA-DA preferentially encodes the reward-predicting cue (lever presentation), whereas NB-ChAT preferentially encodes the reward-motivated action (lever press). (a) Heatmap of GCaMP fluorescence of VTA-DA population for all trials time-locked to sample lever presentation (n=10 recording sites). Each row is a trial, and the trials are sorted by the time of sample lever press relative to sample lever presentation at 0. Orange dotted line is the time of sample lever presentation cue at 0, and the white line is the time of sample lever press action. (b) Same as (a) but in SNc-DA (n=13 recording sites). (c) Same as (a) but in NB-ChAT (n=18 recording sites). (d) Relative contribution of sample lever presentation versus sample lever press in explaining GCaMP fluorescence. Contribution of the predictor of interest is defined as the reduction in explained variance when the predictor was excluded from the full encoding model in Figure 3a. VTA-DA activity preferentially encodes lever presentation cues relative to lever press actions (two-sided paired t-test, p=0.002, n=10 recording sites) (e) same as (d) but in the SNc-DA population (two-sided paired t-test, p=0.22, n=13 recording sites). (f) same as (d) but in the NB-ChAT population (two-sided paired t-test, p=0.02, n=18 recording sites). Note that in contrast to VTA-DA neurons, NB-ChAT activity preferentially encodes lever press actions more than lever presentation cues.

**Supplementary Figure 7.**
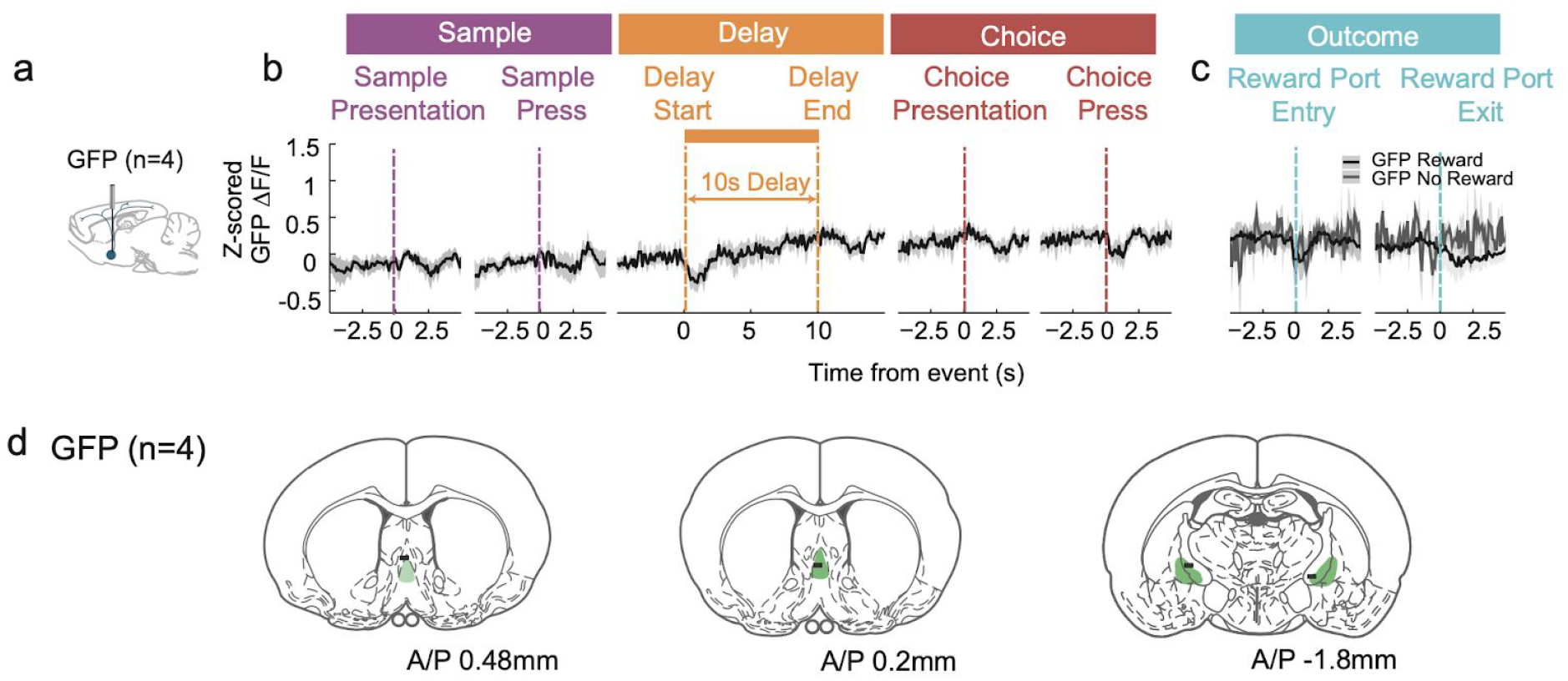
Fluorescence signals from GFP control recording are not modulated by the task. (a) Schematic of fiber photometry recording from GFP control expression sites (b) Z-scored GFP fluorescence from control recordings time-locked to each task event during the sample, delay and choice periods (mean + sem across recordings, n=4 recording sites). Data from all 10s delay trials. (c) Z-scored GFP fluorescence from control recordings during the outcome period, separated by rewarded and unrewarded trials (mean + sem across recording sites, n=4 recording sites). (d) GFP control fiber photometry recording sites (n=4 recording sites). Each line represents the reconstructed location of a fiber tip from histology. Green shaded area is MS and NB.

**Supplementary Figure 8.**
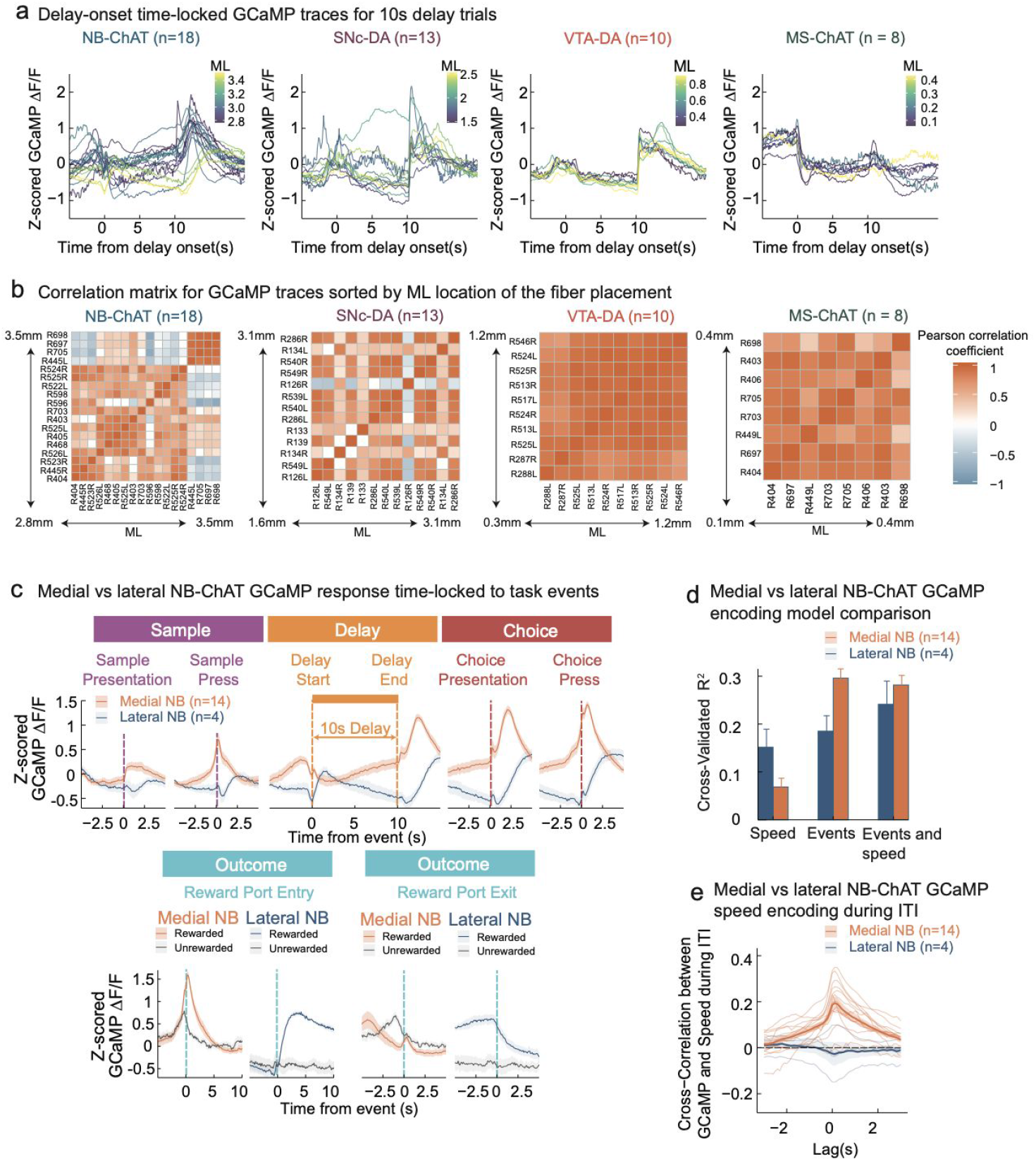
Spatial organization of function across the M-L axis in NB-ChAT. (a) Average Z-scored GCaMP fluorescence from each recording time-locked to delay onset for 10s delay trials (n=10 recording sites for VTA-DA, n=13 recording sites for SNc-DA, n=18 recording sites for NB-ChAT, and n=8 recording sites for MS-ChAT). Color coding denotes M/L distance in mm from midline. (b) Pairwise correlation matrix for GCaMP traces in (a), sorted by ML location of each fiber placement. (c) Medial vs lateral NB-ChAT GCaMP response time-locked to task events. NB-ChAT recording was defined as medial vs lateral based on a cutoff of 3.2mm in ML, based on the division between the two clusters apparent in the cross-correlation matrix in (b) Note that only medial NB-ChAT responds positively to lever press action (sample and choice lever press) and reward. (d) Medial vs lateral NB-ChAT GCaMP encoding model comparisons. Similar to Figure 3b, for each medial vs lateral NB-ChAT group, three encoding models (x-axis) were generated and compared on held-out data: 1) a model with only speed predictors, 2) a model with only task event predictors, 3) the full model with both task event and speed predictors. Lateral NB-ChAT GCaMP fluorescence encodes animal’s speed more strongly than medial NB-ChAT. (e) During the intertrial interval (ITI), the medial NB-ChAT population is positively correlated with speed whereas the lateral NB-ChAT population is not.

**Supplementary Figure 9.**
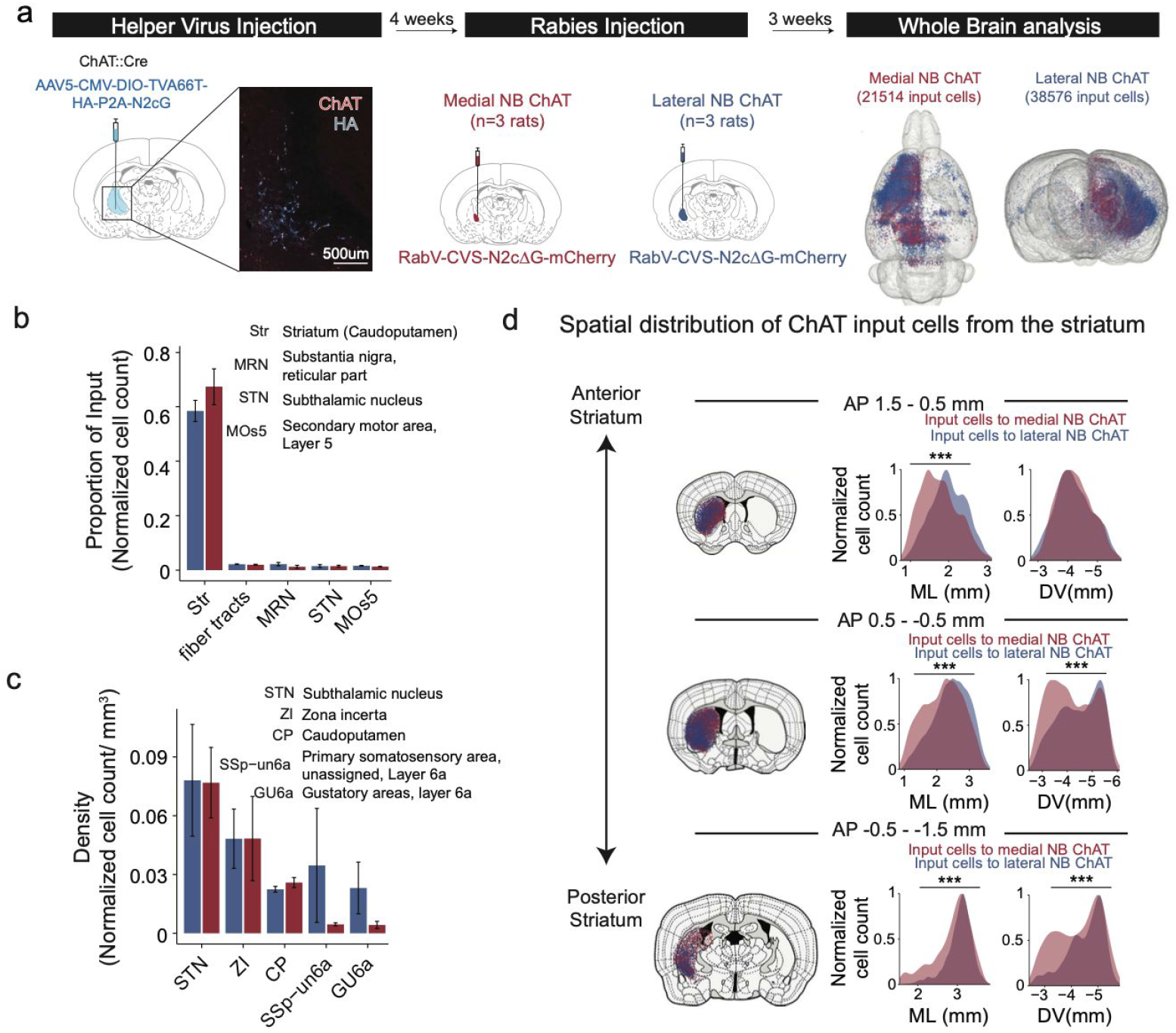
Quantitative whole-brain mapping of inputs to medial vs lateral NB-ChAT. (a) Schematic of experimental design. Cre-dependent helper virus (AAV5-CMV-DIO-TVA66T-HA-P2A-N2cG) is injected into NB of ChAT-Cre rats, limiting its expression to only the NB-ChAT population. 4 weeks later, rabies virus (RabV-CVS-N2c△G-mCherry) is injected into either medial or lateral NB-ChAT, respectively (Reardon et al., 2016). The rabies virus infects all inputs to medial or lateral NB-ChAT and these input neurons are counted and registered into a whole brain (Fürth et al., 2018). (b) Both medial (red bars) and lateral NB-ChAT (blue bars) subregions receive most neuronal inputs from the striatum. y-axis is the proportion of input, defined as the number of structure-specific input neurons normalized by the number of total input neurons in the whole brain. (c) Both medial (red bars) and lateral (blue bars) NB-ChAT subregions receive most neuronal inputs per unit volume from the subthalamic nucleus. y-axis is the proportion of input normalized by the volume of input structure. (d) Medial NB-ChAT receive preferential input from dorsomedial striatum whereas lateral NB-ChAT receive preferential input from dorsolateral striatum. For three subregions of striatum across A/P (1.5 - 0.5mm, 0.5 - −0.5mm, −0.5 - 1.5mm), the spatial distributions of input cells to medial (red curve) and lateral (blue curve) NB-ChAT are compared across M/L and D/V. The input cells to medial NB-ChAT are located more medially than input cells to lateral NB-ChAT in all subregions of the striatum (t-test comparing M/L coordinates of input cells to medial NB-ChAT and M/L coordinates of input cells to lateral NB-ChAT, p<0.001 in all subregions of striatum). The input cells to medial NB-ChAT are located more dorsally than input cells to lateral NB-ChAT in the posterior striatum (t-test comparing D/V coordinates of input cells to medial NB-ChAT and D/V coordinates of input cells to lateral NB-ChAT, p=0.17 in the anterior striatum subregion (A/P 1.5 - −0.5mm); p<0.001 in the posterior striatum subregions (A/P 0.5 - −1.5mm).

**Supplementary Figure 10.**
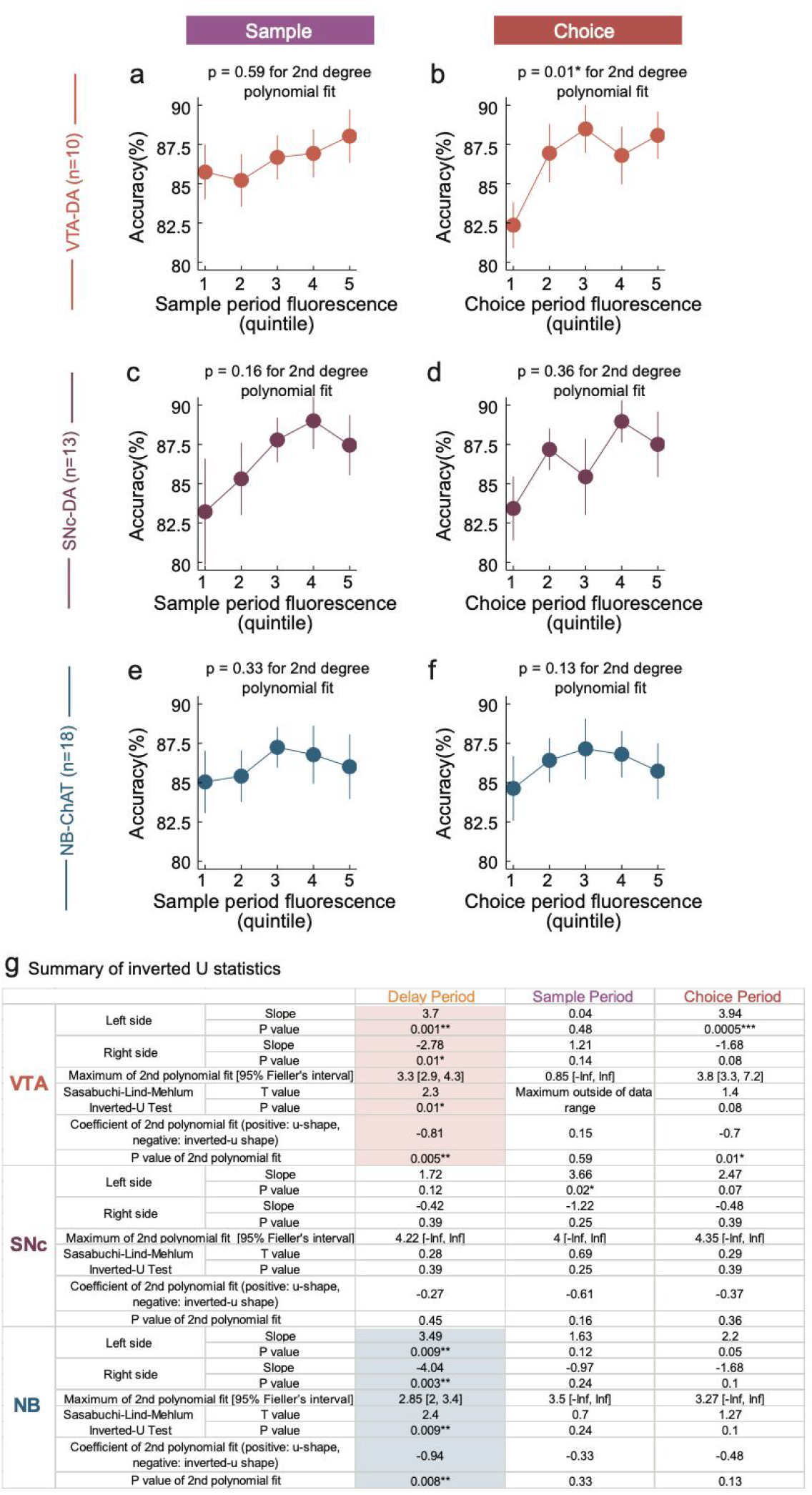
Inverted-U relationship between GCaMP and accuracy is not statistically significant during the sample and choice period. (a) Accuracy relative to sample period fluorescence in VTA-DA population. For each recording, and for each delay length, trials are ranked into quintiles by average sample period GCaMP fluorescence, in order to calculate accuracy for each quintile of fluorescence. Each dot represents accuracy for each sample period quintile, after averaging across delay duration and then across recording sites (mean+sem across recording sites, n=10 recording sites). (b) Accuracy relative to choice period fluorescence in VTA-DA. For each recording, and for each delay length, trials are ranked into quintiles by average choice period GCaMP fluorescence, in order to calculate accuracy for each quintile of fluorescence. Each dot represents accuracy for each choice period quintile, after averaging across delay duration and then across recording sites (mean+sem across recording sites, n=10 recording sites). (c-d) same as (a-b) but in the SNc-DA population (n=13 recording sites). (e-f) same as (a-b) but in NB-ChAT population (n=18 recording sites). (g) Summary of all statistics testing the inverted-U relationship between fluorescence and GCaMP. To meet additional criteria for inverted-U outlined in (Lind and Mehlum 2010), the left side slope of the 2nd polynomial fit has to be significantly positive, the right side slope of the 2nd polynomial fit has to be significantly negative, left and right side should be jointly significant, the maximum of 2nd polynomial fit and its Fieller’s confidence interval have to be within the range of the x-axis, and finally, the coefficient of 2nd polynomial fit has be significantly negative. See Methods for detailed explanation of these additional statistical tests. Only VTA-DA and NB-ChAT delay period fluorescence relationships with accuracy met all aforementioned criteria (highlighted in color).

**Supplementary Figure 11.**
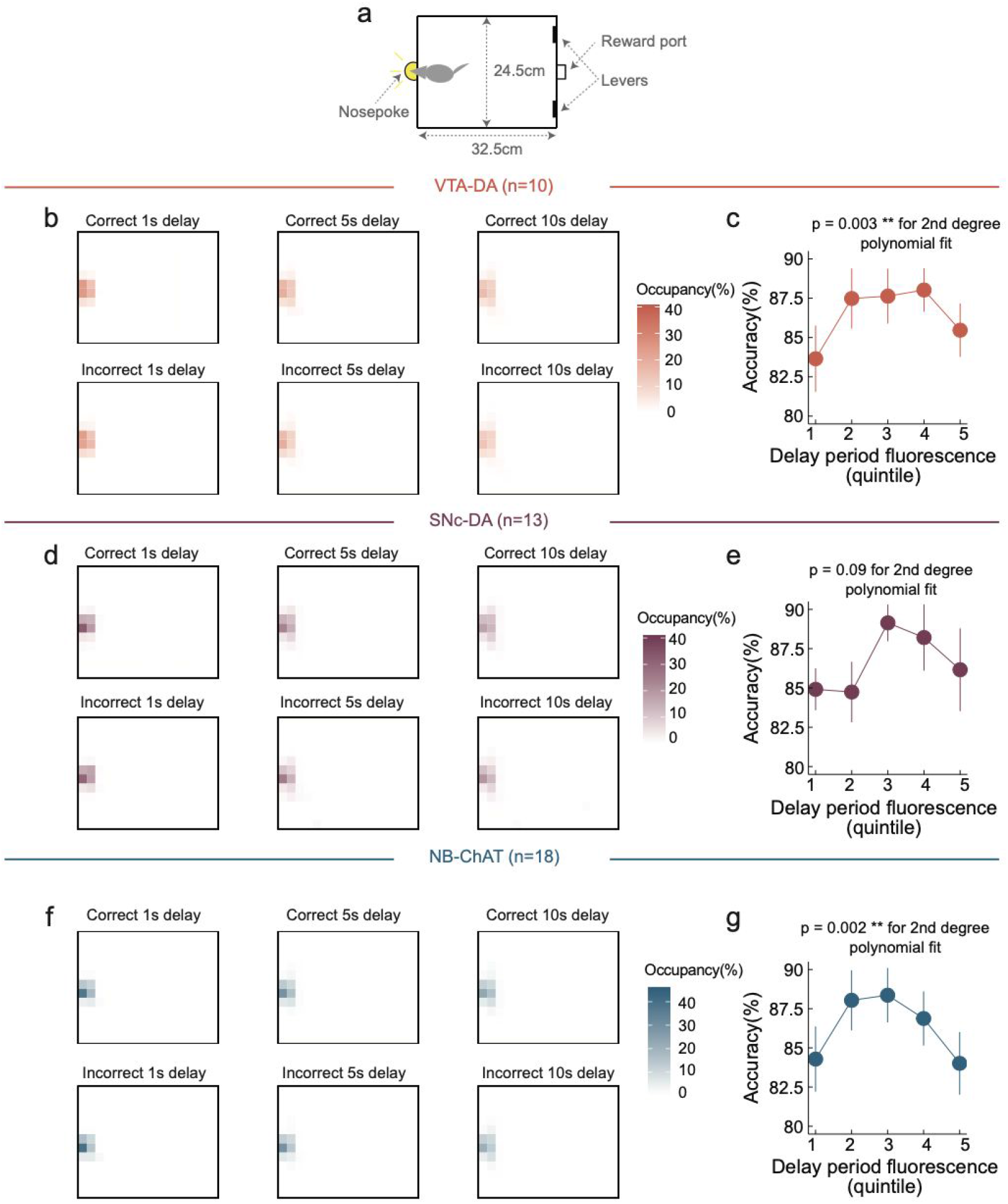
During the delay period, animals spend most of the time near the nosepoke. (a) Schematic of the operant chamber from the top-down view. Nosepoke is located on the left wall, opposite to the levers and reward port. (b) Average position heatmap during all delay durations for correct (top row) and incorrect (bottom row) trials for VTA-DA recordings (n=10 recording sites) (c) Accuracy relative to delay period fluorescence (similar to Figure 3k), using only the subset of data in which the animal’s head was within 10cm of the nosepoke. Each dot represents accuracy averaged across the recording site (mean + sem across recording sites, n = 10 recording sites). (d-e) same as (b-c) but in SNc-DA population (n=13 recording sites). (f-g) same as (b-c) but in NB-ChAT population (n=18 recording sites).

**Supplementary Figure 12.**
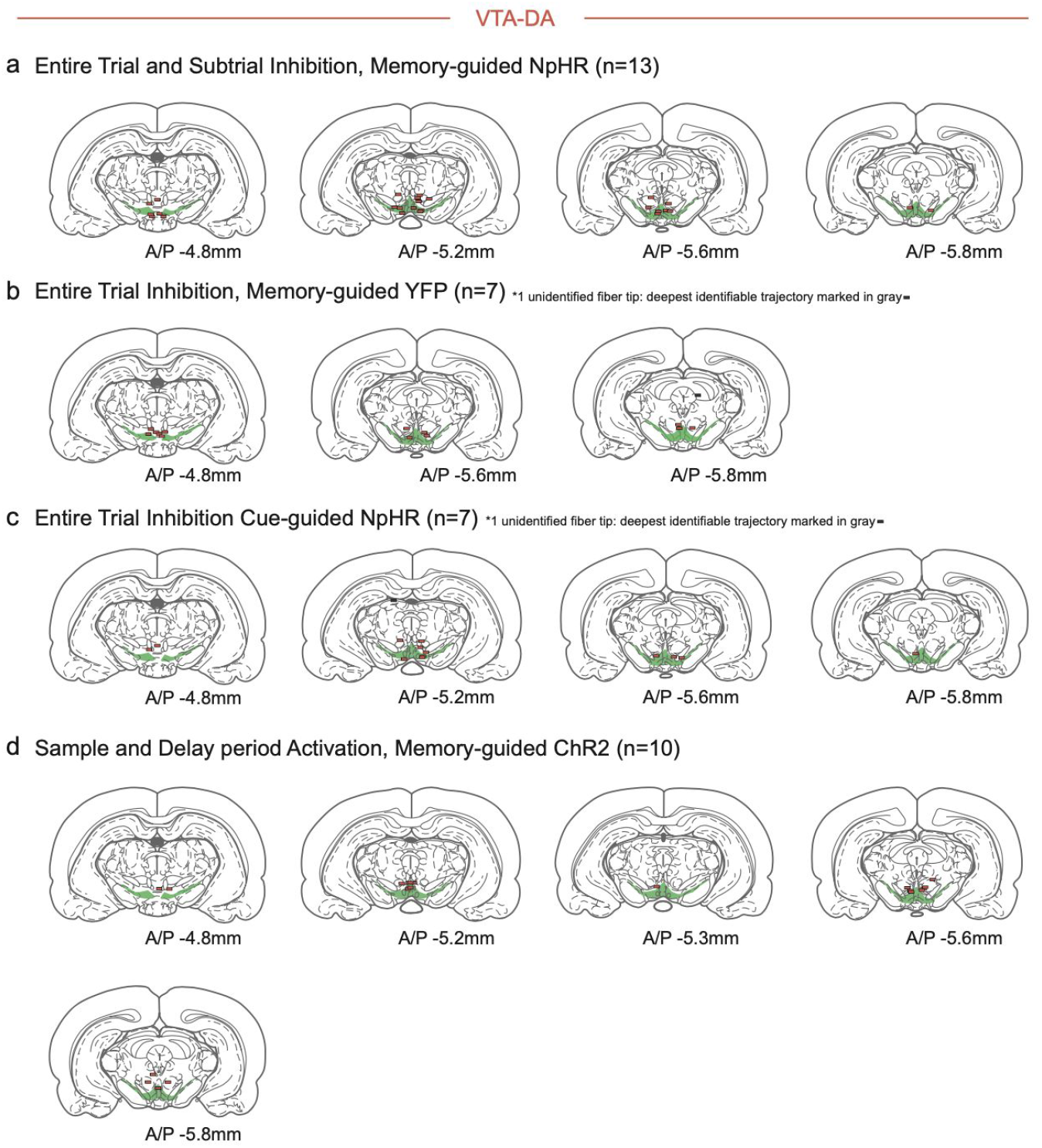
Summary of fiber tip locations for VTA-DA optogenetic manipulations. (a) VTA-DA fiber tip sites from rats used for the entire trial and subtrial inhibition of VTA-DA neurons while performing memory-guided DNMTP task (n=13 rats; data in Figures 5d and 7). Green shaded area is VTA/SNc. (b) VTA-DA optogenetic manipulation sites from rats used for entire trial control illumination of VTA-DA neurons while performing memory-guided DNMTP task (n=7 rats; data in Figure 5e). (c) VTA-DA fiber tip sites from rats used for entire trial inhibition of VTA-DA neurons while performing control cue-guided DNMTP task (n=7 rats; data in Figure 5g). (d) VTA-DA fiber tip sites from rats used for entire trial activation of VTA-DA neurons while performing memory-guided DNMTP task (n=10 rats; data in Figure 8).

**Supplementary Figure 13.**
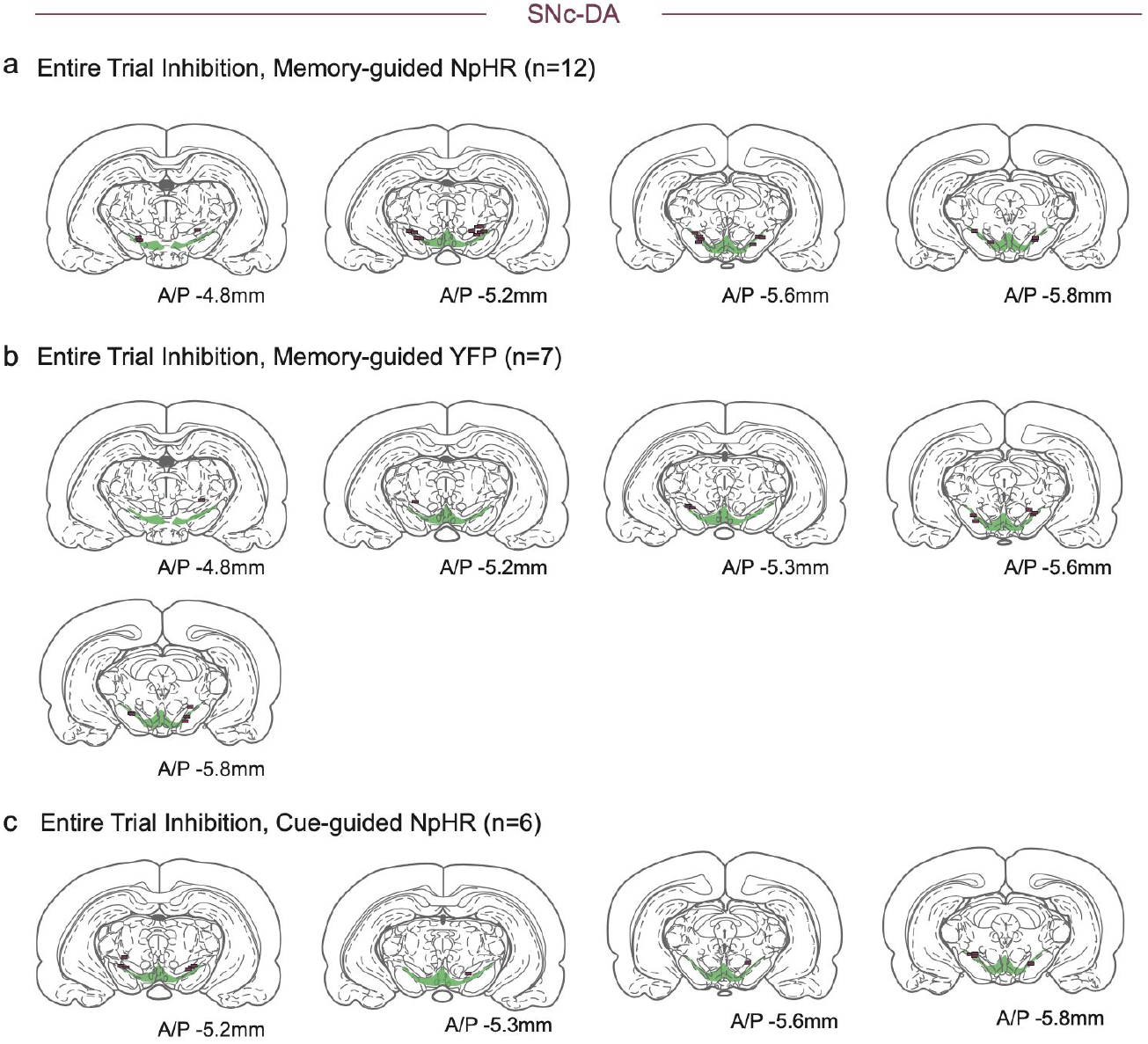
Summary of fiber tip locations for SNc-DA optogenetic manipulations. (a) SNc-DA optogenetic manipulation sites from rats used for entire trial and subtrial inhibition of SNc-DA neurons while performing memory-guided DNMTP task (n=12 rats; data in Figure 5h). Each line represents the reconstructed location of a fiber tip from histology. Green shaded area is VTA/SNc. (b) SNc-DA optogenetic manipulation sites from rats used for entire trial control illumination of SNc-DA neurons while performing memory-guided DNMTP task (n=7 rats; data in Figure 5i). Each line represents the reconstructed location of a fiber tip from histology. Green shaded area is VTA/SNc. (c) SNc-DA optogenetic manipulation sites from rats used for entire trial inhibition of SNc-DA neurons while performing cue-guided DNMTP task (n=6 rats; data in Figure 5k).

**Supplementary Figure 14.**
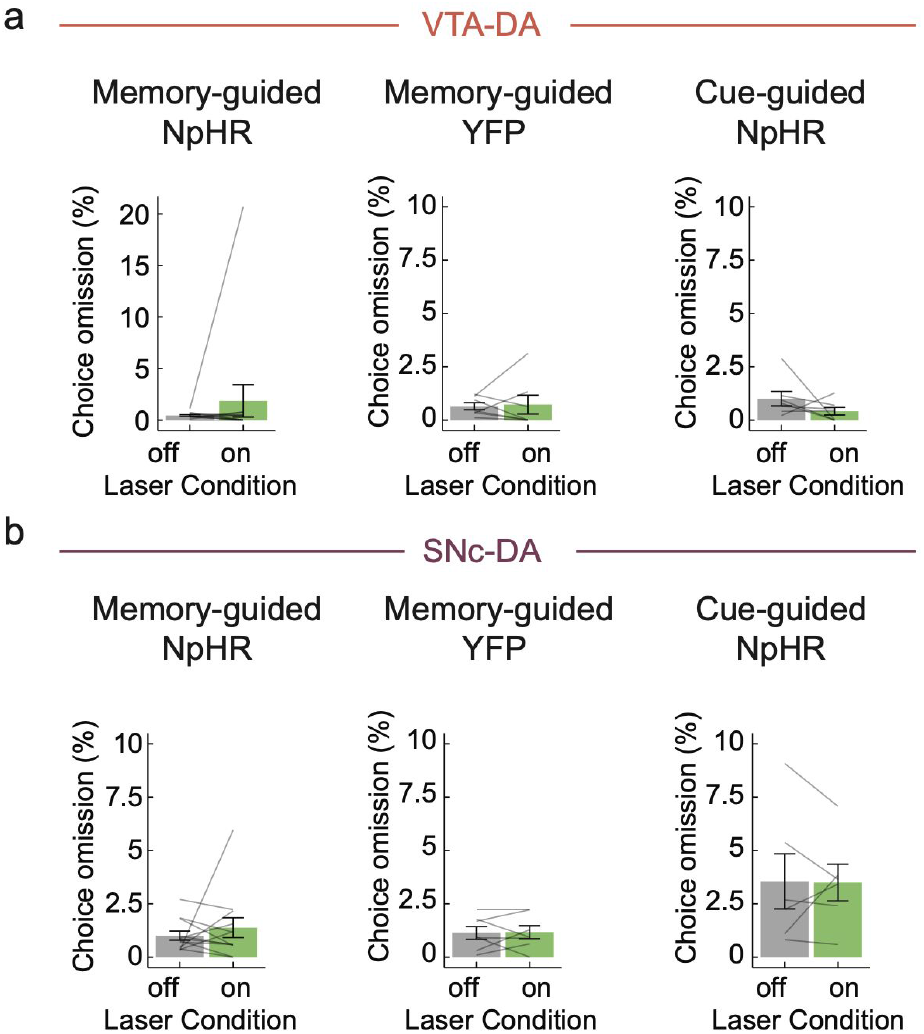
Optogenetic inhibition of VTA-DA and SNc-DA does not affect choice omission rate. (a) VTA-DA optogenetic inhibition does not affect choice omission rate in memory-guided NpHR (left; accuracy data in Figure 5d), memory-guided YFP (middle; accuracy data in Figure 5e), and cue-guided NpHR (right; accuracy data in Figure 5g) experiments. (mixed-effect logistic regression, choice omission/completion predicted based on fixed effects of light, delay, NpHR/YFP opsin group, memory-guided/cue-guided task type, and random effect of individual rat; p=0.23 for light; n=13 rats for memory-guided NpHR, n=7 rats for memory-guided YFP, n=7 rats for cue-guided NpHR) (b) Similarly, SNc-DA optogenetic manipulation does not affect choice omission rate in memory-guided NpHR (left; accuracy data in Figure 5h), memory-guided YFP (middle; accuracy data in Figure 5i), and cue-guided NpHR (right; accuracy data in Figure 5k) experiments (mixed-effect logistic regression, choice omission/completion predicted based on fixed effects of light, delay, NpHR/YFP opsin group, memory-guided/cue-guided task type, and random effect of individual rat; p=0.78 for light; n=12 rats for memory-guided NpHR, n=7 rats for memory-guided YFP, n=6 rats for cue-guided NpHR).

**Supplementary Figure 15.**
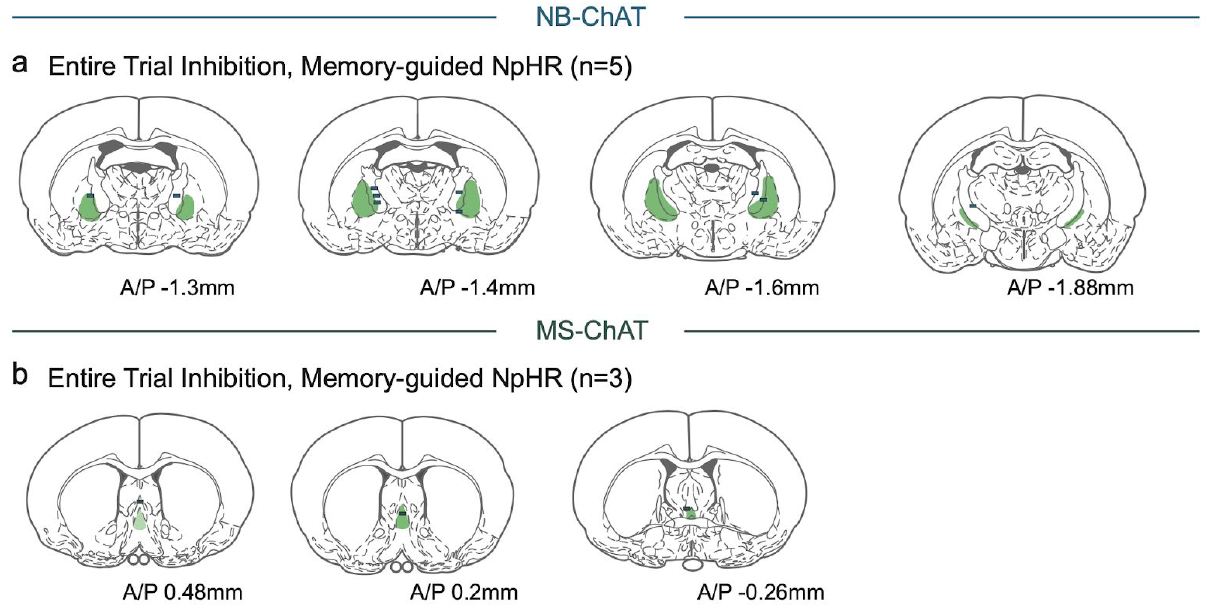
Summary of fiber tip locations for NB-ChAT and MS-ChAT optogenetic manipulations. (a) NB-ChAT optogenetic manipulation locations from rats used for entire trial inhibition of NB-ChAT neurons while performing memory-guided DNMTP task (n=5 rats; data in Figure 6c). Each line represents the reconstructed location of a fiber tip from histology. Green shaded area is NB. (b) same as (a) but from rats used for entire trial inhibition of MS-ChAT neurons while performing memory-guided DNMTP task (n=3 rats; data from Figure 6d).

**Supplementary Figure 16.**
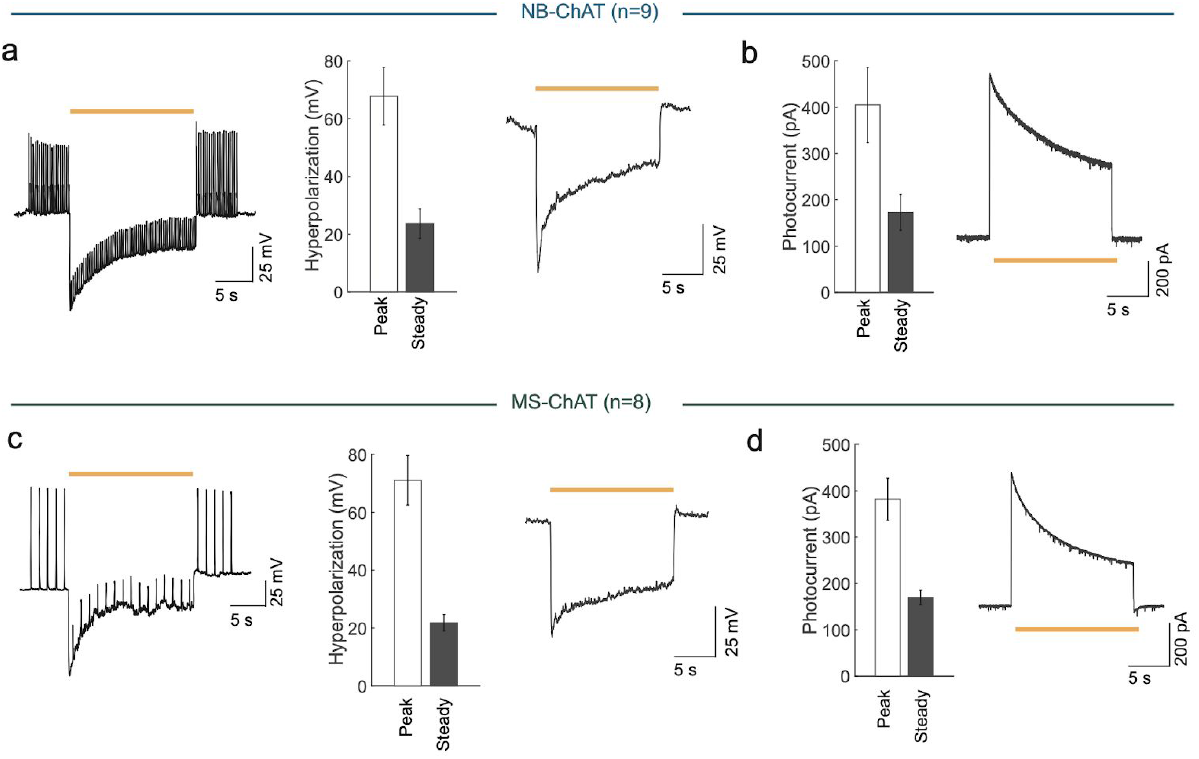
Confirmation of optogenetic inhibition of NB-ChAT and MS-ChAT neurons. (a) Left: Example trace of photoinhibition of spikes generated by current injections (150pA injections, 4 Hz) in NB-ChAT neurons during current clamp recording in brain slices. Middle: Peak and steady-state NpHR-mediated hyperpolarization during current clamp (n=9 neurons; peak hyperpolarization from baseline: 67.8±10.0mV; steady-state hyperpolarization from baseline: 23.7±5.1mV). Right: Example trace of NpHR-mediated hyperpolarization. (b) Left: Peak and steady-state NpHR-mediated photocurrents evoked in NB-ChAT neurons during voltage clamp (n=9 neurons; peak current=404.6±80.9pA; steady-state current=172.8±389.0 pA). Right: Example trace of photocurrent. (c) Left: Example trace of photoinhibition of spikes generated by current injections (200pA injections, 1Hz) in MS-ChAT neurons during current clamp recording. Middle: Peak and steady-state NpHR-mediated hyperpolarization during current clamp (n=8 neurons; peak hyperpolarization from baseline: 71.1±8.5mV; steady-state hyperpolarization from baseline: 21.8±2.9mV). Right: Example trace of NpHR-mediated hyperpolarization. (d) Left: Peak and steady-state NpHR-mediated photocurrents evoked in MS-ChAT neurons during voltage clamp (n=8 neurons; peak current=381.5±45.1 pA; steady-state current= 169.8±16.1pA). Right: Example trace of photocurrent.

## Methods

### Rats

TH::Cre (Horizon TGRA8400) or ChAT::Cre rats (RRRC 658) were maintained on a Long Evans background (Brown et al., 2013; Liu et al., 2016; Witten et al., 2011). A total of 109 rats (108 male and 1 female rats; 36 rats for fiber photometry, 62 rats for optogenetics, 5 rats for slice physiology, and 6 rats for rabies retrograde tracing) weighing > 300g/rat were used for experimentation. At the time of surgery, rats used for fiber photometry experiments were 19+1.01 weeks old, for optogenetics experiments were 18.0±0.77 weeks old, for slice physiology experiments were 14.34±0.03 weeks old, and for rabies retrograde tracing experiments were 12.05±0.39 weeks old. Rats were double-housed, unless they weighed over 500g or had health-related concerns (e.g. fighting). Rats were maintained on a 12-hour light on – 12-hour light off schedule. All surgical and behavioral procedures were performed during the light off cycle.

All experimental procedures were conducted in accordance with the National Institute of Health guidelines and were approved by the Princeton University Institutional Animal Care and Use Committee.

### Delayed non-match to sample (memory-guided) and control cue-guided task

Rats were water-restricted to 80-85% of their *ad-libitum* weight and trained on a delayed non-match to position (DNMTP) spatial short-term memory task in operant chambers (Med-associates; Akhlaghpour et al., 2016; Dunnett et al., 1988). The operant chamber had two retractable levers on the front wall and a nose port on the opposite back wall (Figure 2a). In the DNMTP task, rats were trained to remember the position of the presented sample lever (either right or left) for a delay duration, and report the memory by pressing the “non-match” lever during the choice period. At the beginning of each trial, the sample period was initiated once the sample lever was presented (emerged from the wall) from one of two possible locations - either the right or left position. Upon pressing the sample lever, the lever retracted back into the wall, and the light in the back nose port was illuminated. The delay period started as the rat went to the back wall to poke its nose into the illuminated nose port. The delay period lasted for 1, 5, or 10s (10, 20, 30s or 5, 10, 15s in a subset of experiments shown in Figure 5i and Figure 8) in a randomly interleaved manner, so that the rat did not know when the delay period would end. At the end of the delay period, the nose port lighted up again, and the rats must then make the second nose poke for both levers to extend from the front wall and to begin the choice period. A correct response was to press the lever that did not match the sample lever. A small light in the reward receptacle lit up immediately following the correct lever press, providing a feedback to the rat’s choice as well as signalling the presence of the water reward in the receptacle. Following the feedback light, rats entered the reward receptacle and consumed the water reward. The rats were given up to 15s to press the sample lever and up to 5s to perform a nosepoke in the illuminated nose port and to press the choice lever. All trials were followed by 5s inter-trial interval if the previous trial was correct, and 8s inter-trial interval for previously incorrect or omitted trials.

In the beginning of training, water-deprived rats learned the behavioral sequence of the task to get a reward. Initially, rats spent 1-2 weeks learning a simpler “nose poke-nose poke-lever press” sequence. In the simpler sequence, rats had to make two nose pokes in the back of the chamber, which triggered a random lever to be presented. The pressing of the presented lever led to a drop of water reward. When the rats repeated 100 sequences within an hour of training, they moved onto the more difficult, full sequence, which consisted of “lever press-nose poke-nose poke-lever press”. In this stage, the behavioral sequence was the same as the DNMTP task, but the choice period was modified such that rats needed to simply press the presented lever, instead of making an overt choice between the two levers, as only one choice lever emerged from the wall. At the end of the full sequence, rats were rewarded with a drop of water. The rats learned the full sequence in a few days. Then, the rats were finally introduced to the DNMTP task, in which the two nosepokes were separated by a short time delay (1, 2, 3s), two choice levers were presented, and pressing of the “non-match” to sample lever was rewarded. For the following 3-6 weeks, delays were lengthened (1, 3, 5s, and then 1, 5, 10s) and rats learned the “non-match” to sample rule, improving their performance accuracy (> 80%). In total, the rats received 1-2 months of training.

The cue-guided task served as a control task for DNMTP, as it does not require short-term memory. The task structure was the same as DNMTP with only one difference: the rats were “guided” to the correct choice lever with a cue light directly above the correct lever when the choice levers were presented.

### Surgery

For all surgical procedures, rats were deeply anesthetized in 4-5% isoflurane and placed in a stereotactic setup (Kopf Instruments, Tujunga, CA, USA). After the rats were deeply anesthetized, rats were maintained on 1-2% isoflurane throughout the surgery. The rats received baytril (5mg/kg, i.m.) before surgery and meloxicam (2mg/kg, s.c.) before and 24h after surgery. Rats were allowed a 5 day postoperative recovery period.

#### Fiber photometry experiment

Data in Figures 2-4 are from a series of fiber photometry experiments, which consisted of VTA-DA, SNc-DA, NB-ChAT, MS-ChAT (n=34 rats, 50 recording sites) and control GFP groups (n=2 rats, 4 recording sites).

For the VTA-DA group (n=7 rats, 10 recording sites), 1μL of Cre-dependent GCaMP6f (AAV2/5-CAG-Flex-GCamP6f, Upenn Vector Core, titer: 1.17 × 10^13^ parts/mL or AAV2/5-CAG-DIO-RatOpt-GCaMP6f, PNI Vector Core, titer: 2.30 × 10^13^ parts/mL, (Cameron et al., 2019) was injected into the VTA (A/P: −6.0mm, M/L: 0.8mm, D/V: −8.0mm) of TH::Cre rats.

For the SNc-DA group (n=8 rats, 13 recording sites), 1μL of Cre-dependent GCaMP6f (AAV2/5-CAG-Flex-GCamP6f, Upenn Vector Core, titer: 3.90 × 10^12^ parts/mL or AAV2/5-CAG-DIO-RatOpt-GCaMP6f, PNI Vector Core, titer: 2.30 × 10^13^ parts/mL) was injected into the SNc (A/P: −5.6mm, M/L: 1.7- 2.25mm, D/V: −7.7 - −8.2mm) of TH::Cre rats.

For the NB-ChAT group (n=17 rats, 19 recording sites, note that one recording site was removed from the analysis, see “Encoding models” for details), 1μL of Cre-dependent GCaMP6f (AAV2/5-CAG-Flex-GCamP6f, Upenn Vector Core, titer: 2.34 × 10^12^ parts/mL or AAV2/5-CAG-DIO-RatOpt-GCaMP6f, PNI Vector Core, titer: 2.30 × 10^13^ parts/mL) was injected into the NB (A/P: −1.5mm, M/L: 2.8 - 3.3mm, D/V: −7.0mm) of ChAT::Cre rats.

For the MS-ChAT group (n=7 rats, 8 recording sites), 0.75μL of Cre-dependent GCaMP6f (AAV2/5-CAG-Flex-GCamP6f, Upenn Vector Core, titer: 2.34 × 10^12^ parts/mL) was injected into the MS of ChAT::Cre rats (A/P: +0.5mm, M/L: 0mm, D/V: −7.0mm, 10° angle).

For the control GFP group (n=2 rats, 4 recording sites), 0.75 - 1μL of Cre-dependent GFP virus (AAV2/5-CAG-Flex-eGFP, Upenn Vector Core, titer: 1.81 × 10^12^ parts/mL) was injected to the NB (A/P: −1.5mm, M/L: 3.0mm, D/V: −7.0mm) and MS (A/P: +0.5mm, M/L: 0mm, D/V: −7.0mm) of ChAT::Cre rats.

After the virus injection, a fiber optic cannula (400μm core diameter, low-autofluorescence, MFC_400/430-0.48_10mm_MF2.5_FLT, Doric Lenses) was implanted 0-0.7mm above the injection site. Note that fiber optic cannula implantation into MS and VTA, and virus injection into MS was at 10° angle to divert the superior sagittal sinus.

18 rats contributed two recording sites each (bilaterally or from two different regions), and 18 rats contributed a single recording site each, resulting in a total 54 recording sites from 36 animals.

#### Optogenetics experiment

For the optogenetic inhibition experiment, 1μL of Cre-dependent NpHR (AAV2/5-EF1a-DIO-eNpHR3.0-eYFP, Upenn Vector Core, titer: 1.29 × 10^13^ parts/mL or PNI Vector Core, titer: 1.00× 10^14^ parts/mL) was injected into the SNc and VTA of TH::Cre rats, and NB and MS of ChAT::Cre rats.

For the optogenetic activation experiment, 1μL of cre-dependent ChR2 (AAV2/5-EF1a-DIO-ChR2-eYFP, Upenn Vector Core, titer: 7.70 × 10^12^ parts/mL or PNI Vector Core, titer: 7.0 × 10^14^ parts/mL) was injected into the SNc and VTA of TH::Cre rats.

For the control illumination experiment, 1μL of Cre-dependent YFP virus (AAV2/5-EF1a-DIO-eYFP, PNI Vector Core, titer: 6.0 × 10^13^ parts/ml) was injected into the VTA of TH::Cre rats and SNc of ChAT::Cre rats.

After the virus injection, a fiber optic cannula (300μm core diameter, custom made with MM-FER-2006SS-3300 from Precision fiber products and FT300UMT from Thor labs) was implanted 0-0.7mm above the injection sites. The optogenetic manipulations of VTA, SNc, and NB were bilateral (A/P: −6.0mm, M/L: ±0.8mm, D/V: −8.0mm for VTA; A/P: −5.6mm, M/L: ±1.7 - ±2.25mm, D/V: −7.7 - −8.2mm for SNc; A/P: −1.5mm, M/L: ±2.8 - ±3.3mm, D/V: −7.0mm for NB), and the optogenetic manipulation of MS was unilateral (A/P: +0.5mm, M/L: 0mm, D/V: −7.0mm), since the structure was centrally located in the midline. Also note that the fiber optic cannula implantation into the MS and VTA, and virus injection into the MS was at 10° angle to divert the superior sagittal sinus.

#### Rabies retrograde tracing experiment

In 6 ChAT::Cre rats, 1.5 μL of helper virus (AAV5-CMV-DIO-TVA66T-HA-P2A-N2cΔG, PNI Vector Core, titer: 2.0 × 10^14^ parts/mL) was injected into the NB (A/P: −1.5mm, M/L: 0.75 μL at 2.8mm, 0.75 μl at 3.5mm, D/V: −7.2mm). 4 weeks later, 3 of them were assigned to the medial NB group and received 50, 100, or 200nL of rabies virus injection (RabV-CVS-N2cΔG-mCherry, PNI Vector Core, titer: 2.0 × 10^8^ parts/mL) into the medial NB (A/P: −1.5mm, M/L: 2.8mm, D/V: −7.2mm). The remaining 3 rats were assigned to the lateral NB group and received 50, 100, or 200nL of rabies virus injection into the lateral NB (A/P: −1.5mm, M/L: 3.5mm, D/V: −7.2mm).

#### Ex-vivo slice physiology experiment

In 5 ChAT::Cre rats, 1 μL of Cre-dependent NpHR virus (AAV2/5-EF1a-DIO-eNpHR3.0-eYFP, PNI Vector Core, titer: 2.20 × 10^14^ parts/mL) was injected bilaterally into the NB (A/P: −1.5mm, M/L: ±3.0, D/V: −7.2mm). Additionally, 0.75μL of the same virus was injected into the MS (A/P: +0.5mm, M/L: 0mm, D/V: −7.0mm) at 10° angle to divert the superior sagittal sinus.

### Fiber photometry

We recorded fluorescence through an implanted fiber while the rats were performing the DNMTP task. We excited GCaMP (or GFP in case of control rats) with two different wavelengths: 405nm (intensity at fiber tip: 5-10μW, sinusoidal frequency modulation: 531Hz) and 488nm (intensity at fiber tip: 15-25μW, sinusoidal frequency modulation: 211 Hz) using an LED driver (Thorlabs DC4104). Emission light from GCaMP was collected through the same fiber using a photodetector (Newport, Femtowatt 215), and the analog data was digitized by the TDT system (RZ5D) which served both as a A-D converter and lock-in amplifier. A small head-mounted LED was used to track the rat’s position in the chamber while recording. The position data was simultaneously acquired through the TDT video tracking system (RV2). The timestamps for task events were registered as TTL pulses from the operant chamber into the TDT fiber photometry system through the Med-associates interface connection. Thus, the TDT acquisition system synchronously acquired event time stamps through the Med-associates interface, GCaMP signal through the photodetector, and animal’s head position through the TDT RV2.

#### GCaMP signal processing

With 488nm excitation, the fluorescence of GCaMP is relatively calcium-dependent, but with 405nm excitation, its fluorescence is largely calcium-independent (Akerboom et al., 2012; Tian et al., 2009). When calculating dF/F, we therefore utilized the 405nm channel to calculate the baseline fluorescence in order to account for calcium-independent changes in fluorescence that may be caused by the rats’ movement in our freely moving operant task (Lerner et al., 2015).

The fluorescence signals were acquired at 381Hz and then downsampled to 10Hz using “resample” function in matlab. These downsampled signals were processed according to the following steps:

First, control 405nm signal *S*_*control*_ (*t*) was fit to 488nm GCaMP signal *S*_*GCaMP*_ (*t*) using least-squares regression to calculate the fitted control signal *S*_*fitted*_(*t*) :

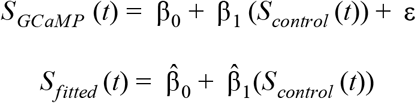

Second, the relative change in fluorescence signal, Δ*F* /*F* (*t*), was calculated using *S*_*GCaMP*_ (*t*) and *S*_*fitted*_(*t*).

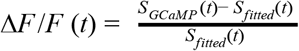

Lastly, Δ*F* /*F* (*t*) was z-scored to facilitate comparison across recording sessions and rats. The mean (*mean*(Δ*F* /*F* (*t*))) and the standard deviation (*std*(Δ*F* /*F* (*t*)) was calculated over each recording session.

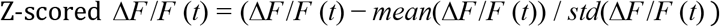

### Encoding models to predict GCaMP from task events and movement

To distinguish the relative contribution of locomotion and task events in predicting the GCaMP signal, we built and compared three encoding models, as shown in Figure 3b. The three models were based on linear regressions, in which the measured GCaMP signal was predicted by the weighted sum of predictors based on task events, animals’ speed in the chamber or the combination of task events and speed.

#### Event predictors

Task event predictors (*E*_*i,j*_) were generated for each type of task events, by convolving a time series of event times (*T*_*i*_, 1 when event occurred, or 0 otherwise) with a 10 degrees-of-freedom spline basis set (*B*_*j*_, where *j* = [1..10]), spanning −1 to +2s around the task events (see (Engelhard et al., 2019; Park et al., 2014). For *i*^*th*^ type of task events and *j*^*th*^ spline basis function, task event predictor (*E*_*i,j*_) is defined as follows:

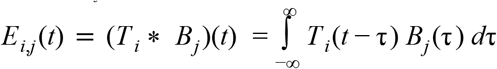

The 10 types of task events consisted of sample lever presentation, sample lever press, delay start, delay end, choice lever presentation, choice lever press, correct reward port entry, correct reward port exit, incorrect reward port entry, and incorrect reward port exit (therefore, *i* = [1..10]). Note the duration of the spline basis set for the reward response (τ) was longer (0-10s) to capture longer reward consumption responses observed in some animals.

The advantage of convolving each event with the spline basis set to generate our predictors is that it allows for a temporal delay in the relationship between neural activity and behavior, while minimizing the number of predictors by assuming smoothness in the response profiles (Engelhard et al., 2019; Park et al., 2014). 10 degrees-of-freedom spline basis set was determined empirically, so that the shape of time-locked GCaMP signal was reasonably captured in the response kernels learned from the model despite the smoothness imposed by the convolution, while minimizing the number of predictors assigned to represent the task event.

#### Speed predictors

Animals’ movement speed was calculated from the tracked x, y position of the rats’ head using a small LED light attached to the fiber photometry tether, close to the rats’ head. The x, y positions were tracked and acquired at 102 Hz. Tracking was lost if the LED light was hidden by the chamber objects (i.e. underneath the lever or too far into the reward consumption inlet) or its reflection on the wall was captured outside of the tracking zone. Missing tracking points were treated as NaN in matlab and R. The tracked x, y position in pixels was converted to centimeters by manually defining the outer edges of the tracked arena, whose dimension was 32.5cm x 24.5cm. The position vectors were iteratively median-filtered three times (with 100ms window) to reduce noise and interpolate missing data from the tracking loss. The Euclidean distance, derived from the change in x, y position, was multiplied by the acquisition frequency to calculate instantaneous speed. The instantaneous speed was then downsampled to 10Hz, using the “resample” function in matlab, to generate the speed predictor.

Speed predictors (*S*^*k*^) were continuous variables which included first, second, and third-degree polynomials of the animal’s speed, to allow flexibility in the relationship between speed and GCaMP.

#### Encoding models

The full encoding model to predict GCaMP (1), a reduced model with only the task event predictors (2), and an alternative reduced model with only the speed predictors (3) are expressed as follows:

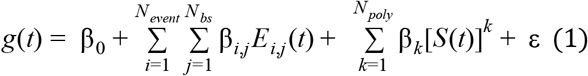

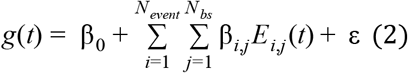

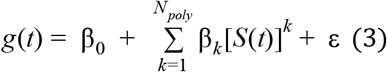

where *g*(*t*) is the predicted GCaMP signal predicted based on task event predictors (*E*_*i,j*_) and/or animal’s speed (*S*^*k*^). Through the linear regression, the model learned β weights (β_0_, β_*i,j*_, and β_*k*_) for the predictors (*E*_*i,j*_ and *S*^*k*^). Parameters in the model include *N*_*event*_, *N*_*bs*_, and *N*_*poly*_, which are defined as the number of types of task events (*T*_*i*_), the degrees of freedom of the spline basis set used for convolution (*B*_*j*_), and the degree of polynomials used to model animal’s speed (*S*^*k*^).

#### Model evaluation using 3-fold cross-validation and R ^2^

To examine the relative contribution of animal’s movement vs. task events predictors, *R*^2^ of the three models, as a measure of model’s predictive power, were calculated and compared (Figure 3b). To generate *R*^2^ of the model, data from each recording site was divided into three folds, in which ⅔ of the data was used to train the model (using the “lm” function in R), and the ⅓ of the data was held-out to test the trained model. After the model was trained, predicted GCaMP from the model was generated using the “predict” function on the predictor matrix of the held-out data. *R*^2^ was then determined by correlating the predicted GCaMP with the recorded GCaMP on a held-out data. This training-testing process was repeated until each fold was used as the held-out data for testing (3-fold cross-validation). The resulting three *R*^2^ for each fold was averaged to create an average rank-deficient fit was not used to calculate average *R*^2^ for each recording site. Note that *R*^2^, since it suggested the data was not sufficient. This resulted in eliminating one NB recording site (1 out of 50 recording sites) from further analysis.

To fit the model with a linear regression, “*lm”* function in R was used (Figure 3 and Supplementary Figure 3a-f). To validate that our model is not overfitting, we also fit the same model using a lasso regression (“glmnet” function in R), which uses regularization to select relevant predictors, thereby reducing the total number of predictors (Supplementary Figure 3g-k).

### Generating event kernels from the model

The response kernel for a type of task event is the component of the neural response that can be specifically attributed to the type of task events in the encoding model. These response kernels learned from the model are reported in Supplementary Figure 3. To generate the response kernels, beta weights (β_*i,j*_) for the task event predictors (*E*_*i,j*_) were learned from regression described above.

For *i*^*th*^ type of task events, response kernels is the weighted (β_*i,j*_) sum of spline basis function *B*_*j*_ (τ) for the task event as follows:

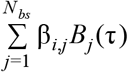

### Inverted-U quantification

To statistically test if there is an inverted-U relationship between fluorescence and accuracy (Figure 4 k,m,o and Supplementary Figure 10a-f), the average accuracy was predicted by a mixed-effect linear regression based on the following predictors: 1st and 2nd degree polynomial of delay period fluorescence quintile, delay period duration, and random effect of individual recording site (implemented with “lmer” function in R). Note that the random effect of individual recording sites allows the model to account for individual differences in average accuracy, while identifying the curve that best fits the entire dataset. The inverted-U was supported by the negative and statistically significant coefficient of the 2nd degree polynomial of delay period fluorescence quintile.

To justify our model selection process, we compared two mixed-effect linear regression models. In the first full model, accuracy was predicted by both the first and second degree polynomial of delay period fluorescence quintile, delay duration, and random effect of individual recording sites. In the second reduced model, everything was the same as the first model, except the second degree polynomial of delay period fluorescence quintile was omitted. Since the second model is nested within the first model, we performed a chi-squared test of the two models to determine if the addition of the second degree polynomial term is justified. In fact, the addition of the second degree polynomial significantly improved the model fit only in the VTA-DA and NB-ChAT group (*X*^2^_6,7_ = 8.22, *p* < 0.001 for VTA-DA, *X*^2^_6,7_ = 7.18, *p* < 0.001 for NB-ChAT), but not in the SNc-DA group (*X*^2^_6,7_ = 0.58, *p* = 0.45 for SNc-DA). The goodness of fit of the selected, full model were 0.58 for VTA-DA, 0.36 for SNc-DA, and 0.45 for NB-ChAT.

To confirm that the statistical significance of the observed inverted-U is not spurious, we repeated the same analysis for shuffled data, which we generated by randomly re-assigning the relationship between accuracy and delay period fluorescence. As expected, this shuffling procedure eliminated the significance of the inverted-U (for shuffled delay period data: p=1.0 for VTA-DA shuffled delay period fluorescence, p=0.64 SNc-DA shuffled delay period fluorescence, p = 0.45 for NB-ChAT shuffled delay period fluorescence).

We further validated the inverted-U by incorporating an additional set of statistical tests, based on (Lind and Mehlum, 2010). These results are summarized in Supplementary Figure 10g. They recommend that in addition to the 2nd degree polynomial p-value described above, an inverted-U should be confirmed through: i) significance of the positive slope on the lower data range, ii) significance of negative slope on the upper data range, iii) joint significance of the left and right side slope, and iv) checking that the maximum of the inverted-U and its confidence interval fall within the x-range of the data. Given the inverted-U equation (y=*α*+*β*x+*γ*x^2^), the significance of positive and negative slopes was computed from one-sided t-test for inequalities *β*+2*γ*x_l_ < 0 and *β*+2*γ*x_h_ > 0, where x_l_ and x_h_ were the minimum and maximum of the data range (in this case, the 1st and 5th quintile of fluorescence). The joint significance of the two slopes was tested from the composite hypotheses of the inequalities (*β*+2*γ*x_l_ < 0 ∪ *β*+2*γ*x_h_ > 0, intersection-union test). Fieller’s confidence interval around the maximum of the inverted-U exploits the fact that the maximum can be expressed as a ratio of two normally distributed estimates 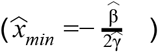 and that the distribution of such ratio is also normal (Fieller, 1940). Accordingly, Fieller’s (1-*α*) confidence interval of the ratio can be computed by finding a set of *θ* values:

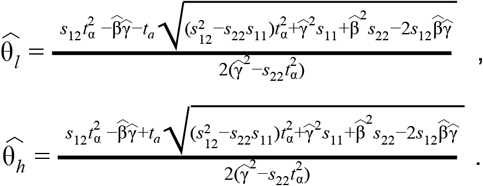

where *s*_11_, *s*_22_ and *s*_12_ are the estimated variances of *β* and *γ*, and the covariance between them respectively, and *t*_*a*_ is the t-statistic from the aforementioned intersection-union test of the composite hypotheses of inequalities, *β*+2*γ*x_l_ < 0 ∪ *β*+2*γ*x_h_ > 0. These statistics were computed using the Stata module provided with the paper (https://econpapers.repec.org/software/bocbocode/s456874.htm)

### Relative contribution of lever presentation vs lever press

To compare the relative contribution of reward-predicting cues and reward-motivated actions in predicting GCaMP fluorescence (Supplementary Figure 6), we quantified the reduction in variance explained when the predictor of interest was removed from the encoding model.

First, we compared the full model (as described earlier in the “Encoding models to predict GCaMP from task events and movement” section) with a cue-reduced model. The cue-reduced model was the same as the full model, except the “sample lever presentation” predictors (10 basis set predictors for the “sample lever presentation” event) were removed from the predictor matrix. The data was fit again to the cue-reduced model, using the “lm” function and 3-fold cross-validation. The contribution of the sample cue predictors (*C*_*cue*_) was defined as the reduction in the explained variance, *R*^2^, of the reduced model compared to the full model (Engelhard et al., 2019; Lovett-Barron et al., 2019; Musall et al., 2019):

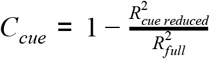

We similarly compared the full model with an action-reduced model by removing the “sample lever press” predictors from the predictor matrix and calculating the contribution of sample lever press predictors (*C*_*action*_):

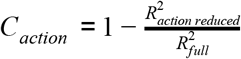

Note that we removed sample presentation and sample press (and not choice presentation or choice lever press) to derive the cue-reduced and action-reduced models. This is because choice press coincided with the light cue for reward in our task design, thus we were unable to cleanly dissociate the reward cue from choice lever press action.

Finally, the relative contribution of the predictor for each recording site was calculated as a percentage over the combined contribution of cue and action.

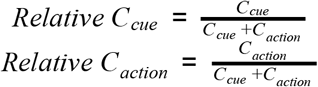

To statistically compare the relative contribution of cues and actions to the explained variance, we performed pairwise t-tests across the VTA-DA, SNc-DA, and NB-ChAT recording sites.

### Immunohistochemistry

Rats were deeply anesthetized using euthasol (2mg/kg, i.p) and transcardially perfused first with phosphate-buffered saline (PBS), and then with 4% paraformaldehyde (PFA) in PBS. Brains were collected and post-fixed in 4% PFA overnight. The brains were then placed in 30% sucrose in PBS solution for 2-5 days at 4°C. Frozen brains were cut into 40-50μm thick coronal sections using a cryostat.

One-third of the coronal sections near the target location were directly mounted from the cryostat and cover-slipped with a mounting solution (fluoromount-G with DAPI, Southern Biotech) to obtain accurate fiber location and to confirm virus expression without any staining. These images were taken using a microscope (Nikon Ti2000E or Leica M205FA) or whole slide scanner (Hamamatsu Nanozoomer S60).

Another one-third of sections were stained for TH or ChAT, to observe co-localization with GCaMP, NpHR, or ChR2. These sections were placed in a blocking buffer (2% normal donkey serum and 1% bovine serum albumin in PBST; Sigma A7906-100G) for 30min. Then for TH staining, sections were incubated overnight at 4°C in solution containing the primary antibody for tyrosine hydroxylase (Chicken-TH, 1:500 or 1:1000 dilutions, Aves lab TYH). For ChAT staining, sections were incubated for two days at 4°C in solution containing the primary antibody for choline acetyltransferase (Goat-ChAT, 1:100 dilution, Millipore AB144P). When enhancement of GCaMP, NpHR, and ChR2 signals was necessary, primary antibody for GFP was used (Rabbit-GPF, 1:1000 dilution, Molecular Probes G10362). Sections were then washed with PBS for 30min, and incubated overnight at 4°C in Alexa Fluor 647 or Cy3 (Donkey anti-Chicken-Cy3, 1:1000 dilution, Jackson ImmunoResearch, 703-165-155 or Donkey anti-Goat-647, 1:1000 dilution, Jackson ImmunoResearch, 705-605-147) and Alexa Fluor 488 (Donkey anti-Rabbit-488, 1:1000 dilution, Jackson ImmunoResearch, 711-545-152). After PBS washes, sections were mounted in a mounting solution (fluoromount-G with DAPI, Southern Biotech). To confirm colocalization, cellular resolution images were taken using a confocal microscope (Leica TCS SP8).

### Reconstruction of fiber placement from histology sections

Fiber tip locations of the fiber photometry recording sites (Supplementary Figures 1, 8, 9) were reconstructed from the histology of coronal brain sections referencing the Paxinos Rat Atlas (Paxinos and Watson, 6^th^ edition). A/P position of the fiber tip was approximated from the section with the deepest fiber track.

In the section with the deepest fiber tip location, M/L position of the fiber tip was carefully reconstructed by normalizing the measured M/L distance of the fiber tip to the reference M/L distance, and scaling that ratio to match the Paxinos Rat Atlas. These normalization-scaling steps effectively registered the measured M/L position into the Paxinos atlas, accounting for individual tissue shrinkage in each brain during histology. Reference distance utilized well-defined “landmarks” in the tissue, such as the distance from the midline to the outermost edge of the tissue (i.e. longest M/L). Then, we derived the atlas-referenced M/L distance of the fiber tip by equating the ratio of measured M/L distances of fiber tip and reference mark to the ratio of atlas-referenced M/L distances of the fiber tip and the reference landmark, and solving for the atlas-referenced M/L distance of the fiber tip.

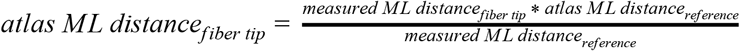

The D/V position of the fiber tip was also derived similarly by referencing the distance of well-known “landmarks” along the D/V (e.g. D/V distance from the top to bottom of the tissue along the midline).

### Quantification of opsin expression level

We quantified the fluorescence intensity as a measure of opsin expression level and correlated it with light-induced accuracy impairment (Figure 5f and 5j). To do so, we collected the tissue with the deepest fiber track and imaged them under the same setting using a Leica M205FA microscope. Using Leica LAS X software, we manually drew the outline of fluorescent areas (around VTA/SNc region). The fluorescence intensity inside the fluorescent area was measured and then normalized by the fluorescence intensity outside the fluorescent area.

### Optogenetic experiment

About 6-7 weeks post virus injection (AAV2/5-EF1a-DIO-NpHR-eYFP in experimental group, AAV2/5-EF1a-DIO-eYFP in control group, for detail, see Methods, Surgery, Optogenetics experiment), rats were tested in the entire-trial inhibition experiment (Figure 5d-e, 5h-i). In randomly-selected 20% of all trials, rats received green light bilaterally throughout the sample, delay, and choice periods (532nm continuous illumination, 5-6mW intensity at the fiber tip) in SNc and VTA for 5 sessions. Rats in the VTA and SNc groups performed ~242 trials/session and ~212 trials/session on average respectively.

For entire-trial inhibition experiment in the NB and MS (Figure 6c-d), rats received green light bilaterally throughout the sample, delay, and choice periods (532nm continuous illumination, 5-6mW intensity at the fiber tip) on NB and MS for 2 sessions in 15% of all trials. Each test session was 1.5 hour long and interleaved with a day where rats performed the task without illumination in order to reduce behavioral adaptation to the manipulation. Rats in the NB and MS groups performed ~294 trials/session and ~308 trials/session on average respectively.

Rats expressing NpHR in VTA were then used for the sub-trial inhibition experiment (Figure 7b-d). Rats received green light (532nm continuous illumination, 5-6mW intensity at the fiber tip) in SNc or VTA in a randomly-selected 30% of all trials for 10 testing sessions, with each test session interleaved with a day where rats performed the task without testing.The laser-on trials were randomly and equally distributed into sample light-on trials (10% of total), delay light-on trials (10% of total), and choice light-on trials (10% of total). Rats performed an average of ~238 trials/session.

A subset of rats from the aforementioned entire-trial inhibition experiments (n=5 for VTA, n=3 for SNc) and additional rats (n=2 for VTA, n=3 for SNc) were trained on the cue-guided task to use cue light to guide their choice (Figure 5g, 5k). As they quickly learned the new rule (in ~2 weeks), they reached >95% average accuracy in all delays (and delay-dependence accuracy impairment dissipated in re-trained rats). These rats received entire-trial inhibition using the same parameter (20% 532nm continuous green light-on trials, 5-6mW, 5 sessions) from the DNMTP entire-trial inhibition experiment. Rats in the VTA and SNc groups performed ~240 trials/session and ~225 trials/session on average respectively.

For the ChR2 experiments, a separate cohort of rats were injected with DIO-ChR2-eYFP in the VTA and tested 6-7 weeks post-injection. For sample period activation experiment (Figure 8d), rats received pulsed blue light in VTA when the sample lever was presented (447nm, 5ms pulse duration, 1 burst of 5 pulses at the sample presentation, ~15mW intensity at the fiber tip). For delay period activation experiment (Figure 8f, 8h), rats received pulsed blue light in the VTA during the delay period (447nm, 5ms pulse duration 20Hz burst per second of 5 pulses or 1pulse per second, ~15mW intensity at the fiber tip). Stimulation took place on a randomly selected 20% of all trials for a total of 5 stimulation sessions, interleaved with nonstimulation sessions. Rats performed on average ~187 trials/session.

### *Ex vivo* electrophysiology recordings to confirm inhibition of MS and NB ChAT cells

To test the efficacy of optogenetic inhibition in MS-ChAT and MS-ChAT cells, we performed ex vivo electrophysiology in ChAT-Cre rats (Supplementary Figure 16). Coronal slices containing the MS or NB were prepared from 5 month old male ChAT-Cre rats 4 weeks after injecting with DIO-NpHR virus. Rats were deeply anesthetized with an intraperitoneal injection of euthasol (2mg/kg, ip) and decapitated. After extraction, the brain was immersed in ice-cold carbogenated N-methyl-D-glucamine (NMDG) artificial cerebrospinal fluid (ACSF) (92 mM NMDG, 2.5 mM KCl, 1.25 mM NaH2PO4, 30 mM NaHCO3, 20 mM HEPES, 25 mM glucose, 2 mM thiourea, 5 mM Na-ascorbate, 3 mM Na-pyruvate, 0.5 mM CaCl2·4H2O, 10 mM MgSO4·7H2O and 12 mM N-acetyl-L-cysteine) for 3 min. Afterwards, coronal slices (300 μm) were sectioned using a vibratome (VT1200s, Leica) and then incubated in NMDG ACSF at 34 °C for 12-14 min. Slices were then transferred into a holding solution of HEPES ACSF (92mM NaCl, 2.5mM KCl, 1.25mM NaH2PO4, 30mM NaHCO3, 20mM HEPES, 25mM glucose, 2mM thiourea, 5mM Na-ascorbate, 3mM Na-pyruvate, 2mM CaCl2·4H2O, 2 mM MgSO4·7H2O and 12mM N-acetyl-L-cysteine, bubbled at room temperature with 95% O2, 5% CO2) for at least 45 min until recordings were performed. Whole cell recordings were performed using a Multiclamp 700B (Molecular Devices, Sunnyvale, CA) using pipettes with a resistance of 4-7MOhm filled with a potassium-based internal solution containing 120mM potassium gluconate, 0.2mM EGTA, 10mM HEPES, 5mM NaCl, 1mM MgCl2, 2mM Mg-ATP and 0.3mM NA-GTP, with the pH adjusted to 7.2 with KOH. ChAT neurons were identified for recordings based on YFP expression. Photostimulation parameters were 586nm and 0.034-0.053mW/mm^2^. Neurons were held at −70mV during photocurrent measurements. Baseline potential was calculated as the mean potential over a 1s period just prior to stimulation. Peak hyperpolarization was calculated as the largest hyperpolarization relative to baseline potential. Steady-state hyperpolarization was calculated as the mean hyperpolarization during the last 1s of stimulation. Peak and steady-state photocurrents were calculated using the same time intervals. To confirm the ability of photocurrents to eliminate action potentials in MS ChAT cells, action potentials were induced by a positive current injection (200pA, 25ms pulse duration, 1Hz). Action potentials in NB ChAT cells were induced by a positive current injection (150 pA, 50ms pulse duration, 4Hz). Stimulation frequencies were chosen based on published in vivo firing frequencies of either cell population (Hedrick and Waters, 2010; Simon et al., 2006).

### Rabies tracing and whole-brain quangification

To analyze input cells to medial and lateral subregions of the NB-ChAT population, we injected Cre-dependent helper virus and rabies virus into the NB of ChAT::Cre rats (for detail, see Methods, Surgery, Rabies retrograde tracing experiment; Supplementary Figure 9; Reardon et al., 2016). 3 weeks post surgery, rats (n=6 rats, 3 rats in each medial and lateral NB-ChAT groups) were perfused and their brains were extracted for histology (for detail, see Immunohistochemistry). Brain sections covering the entire brain (approximate AP range from +4 - −9mm) in 100μm spacing were mounted and cover-slipped with a mounting solution, then imaged using a whole slide scanner (Hamamatsu Nanozoomer S60). These images (Raw 16-bit TIFF) of brain sections were analyzed using a published platform, “WholeBrain” (Fürth et al., 2018).

The analysis of the brain sections consisted of three steps - registration to Allen brain atlas, detection of input cells, and final registration to Waxholm Space atlas of the Sprague Dawley rat brain. First, we visually identified the corresponding mouse A/P coordinate of all rat brain sections, referencing Openbrainmap (http://openbrainmap.org). Then each imaged section of the rat brain were registered into the Allen brain atlas of the same A/P coordinate, using the “registration” function from “Wholebrain” package in R. Once the imaged section was registered, mCherry-labeled cells (input cells infected with RabV-CVS-N2cΔG-mCherry virus) in the images were automatically detected using the “segment” function from “Wholebrain” package in R, with visual inspection to detect outliers and manually correct when deemed necessary. When the registration and detection steps are over, “Wholebrain” creates a data frame containing information on all counted mCherry-labeled cells, their location (A/P, M/L, D/V) in Allen brain atlas, and the brain ontology they belong. Finally, an additional registration process converted the mouse brain coordinates of the detected input cells into the rat brain coordinates using the new “map.to.rat” function (WholeBrain v. version 0.1.36).

